# Remodeling of the immune-metabolic landscape triggered by long-term high-altitude exposure

**DOI:** 10.1101/2025.07.21.665899

**Authors:** Jiaqi Wang, Yutong Dong, Ruoyi Xue, Yi Huang, Wubin Yang, Chen Zhang, Yangkai Zhang, Fengsheng Wang, Ran Yang, Jiangjun Wang, Meng Yu, Yixiao Xu, Manying Guo, Yanping Tian, Rui Jian, Junlei Zhang, Yan Ruan, Yan Hu

## Abstract

Growing evidence indicates that immunological and metabolic outcomes are key mediators of long-term high-altitude exposure (LTHAE) adaption, but the underlying mechanisms remain poorly understood. This study employs plasma metabolomics and peripheral blood single-cell transcriptomic sequencing to analyze the metabolic and immune dynamic regulation in 46 young male lowlanders following a 90-day adaptation period at high altitude. Single-cell analysis shows a pattern of “innate immune activation and adaptive immune suppression” under LTHAE, characterized by facilitated maturation of neutrophils, enhanced cytotoxicity of CD56^dim^ NK cells, and increased immune responsiveness of cDC2 and pDC, while inhibited maturation of plasmablasts and suppressed immune responsiveness of CD8□TEM and CD4^+^ T cells. Plasma metabolic analysis reveals significant alterations, involving enhanced steroid hormone synthesis, unsaturated fatty acid and amino acid metabolism under LTHAE, which in turn are associated with immune remodeling. Moreover, transcriptomic-metabolic integration analysis indicates the molecular mechanisms of enhanced aerobic oxidation efficiency under LTHAE. Collectively, these findings provide integrated insights into immune-metabolic landscape remodeling and suggest potential mutual regulatory relationship between immune and metabolic state following LTHAE, offering a molecular foundation for high-altitude adaptation research.

## Introduction

In recent years, with the rapid growth of tourism, mountaineering expeditions and long-term residents in plateau areas, the comprehensive effects of long-term high-altitude exposure (LTHAE) (defined as continuous residence at altitudes ≥1500 m for over one month) on human health have become an important issue in the field of public health^1–4^. The impact of the plateau environment, characterized by low pressure, low oxygen levels, and strong ultraviolet radiation, on human health exhibits complex and cumulative features. While lowlanders exhibit immediate adaptive responses to acute hypoxia—including hypoxic ventilatory response (HVR), tachycardia, and sympathetic activation—these transient physiological responses typically resolve within 72 hours of descent^5–7^. In contrast, LTHAE can induce persistent pathophysiological adaptations such as polycythemia in the bone marrow, pulmonary vascular remodeling, and neuroendocrine compensation^8^. Prolonged activation of these compensatory mechanisms may progress to irreversible damage, evidenced by epidemiological data showing a 5-18% prevalence of high-altitude polycythemia among residents, alongside elevated risks of high-altitude pulmonary hypertension and cerebral edema^9–11^. Furthermore, chronic exposure correlates with systemic health deterioration, including renal dysfunction, and neurocognitive decline, severely compromising quality of life and occupational capacity^1, 12^. Clarifying the pathophysiological, immune and cellular biological basis of LTHAE mediating multi-organ damage will provide a key theoretical basis for achieving early warning and targeted intervention of high-altitude related diseases.

In the face of complex and diverse environmental stressors, such as extreme temperature, pathogen invasion, radiation or hypoxia exposure, the body maintains homeostasis by activating a series of physiological adaptive mechanisms. Immune regulation and metabolic reprogramming, as two core regulatory networks, coordinate energy allocation and immune defense through dynamic interaction^13, 14^. The hypobaric hypoxia characteristic of high-altitude environments acts as a prototypical systemic stressor, triggering abrupt declines in blood oxygen partial pressure, mitochondrial oxidative phosphorylation impairment, and reactive oxygen species (ROS) bursts^15^. These cascading effects drive systemic metabolic dysregulation and immune imbalance, exacerbating oxidative damage, microvascular dysfunction, and tissue fibrosis—pathological hallmarks underlying high-altitude pulmonary edema, cerebral edema, and multiorgan failure^16^. The metabolic characteristics of plateau residents reveal that enhancing the activity of glycolytic pathway and reducing oxidative stress level are important strategies to adapt to hypoxia^17^. Previous studies have found that acute high-altitude exposure can cause dynamic changes in peripheral immune cell subsets and characteristic metabolite profiles^18, 19^. Indeed, there is a close interaction between the metabolic remodeling induced by high-altitude and the dynamic regulation of the immune system. Metabolic remodeling not only supports cellular energy demands but also influences the differentiation and function of immune cell subsets through specific metabolic intermediates^14^. However, the metabolic and immune characteristics of lowlanders following LTHAE, especially at the single-cell level, remain incompletely understood, and the correlation between these characteristics has not been elucidated.

In this study, we systematically analyzed the immune cell remodeling and metabolic characteristics in lowlanders following LTHAE using peripheral blood single-cell transcriptome sequencing and plasma metabolomics. This study aims to provide theoretical support for deeper understanding of immune homeostasis regulation during high-altitude adaptation and to explore molecular targets for intervening in high-altitude-related diseases, potentially contributing to precision prevention strategies for mountain sickness.

## Results

### 1. Experimental design and physiological parameters of the subjects

A total of 46 young male subjects (age 22.0±1.6 years, BMI 22.7±1.5 kg/m²) were enrolled in this study. In a low-altitude (LA) environment (altitude 200 m), physiological assessments were conducted on the 46 subjects at rest, and fasting venous blood samples were collected from them for routine blood tests, biochemical analysis, and plasma metabolomics. Among these, the blood samples from 4 subjects were used for peripheral blood single-cell sequencing. After the subjects were stationed in high-altitude (HA) environment of Lhasa, Tibet (altitude 3650 m) and adapted to the plateau for 90 days, the aforementioned indicators were retested (Figure 1). From plain to plateau, oxygen saturation of pulse oximetry (SpO□) decreased significantly from 99.2±0.5% to 86.7±2.5%, systolic blood pressure decreased from 116.0±10.5 mmHg to 105.8±11.9 mmHg. Meanwhile, the heart rate increased from 74.2±9.8 beats/min to 91.0±14.4 beats/min in a compensatory manner, suggesting that the cardiovascular system alleviated tissue hypoxia by increasing cardiac output. The red blood cell count increased from 4.9±0.3×10¹²/L to 5.8±0.3×10¹²/L, hemoglobin levels rose from 158.0±54.4 g/L to 172.0±8.3 g/L, and platelet count went up from 252.9±46.1×10□/L to 273.8±48.7×10□/L, all of which are consistent with the characteristics of bone marrow hyperplasia induced by long-term hypoxia^20^. The total white blood cell count increased from 6.0±1.2×10□/L to 7.0±1.6×10□/L, with significant rises in neutrophils, lymphocytes, and monocytes, while eosinophils and basophils decreased, indicating that LTHAE triggered the reconstitution of circulating immune cells. Creatine kinase (CK) and its isoenzyme (CK-MB) increased to 1692.5±5863.0 U/L (baseline 203.6±248.7) and 38.7±56.4 U/L (baseline 12.6± 5.5), respectively, suggesting that high-altitude exposure may cause damage to skeletal muscle and myocardial mitochondrial function (Supplementary Table 1-2).

**Figure 1.**
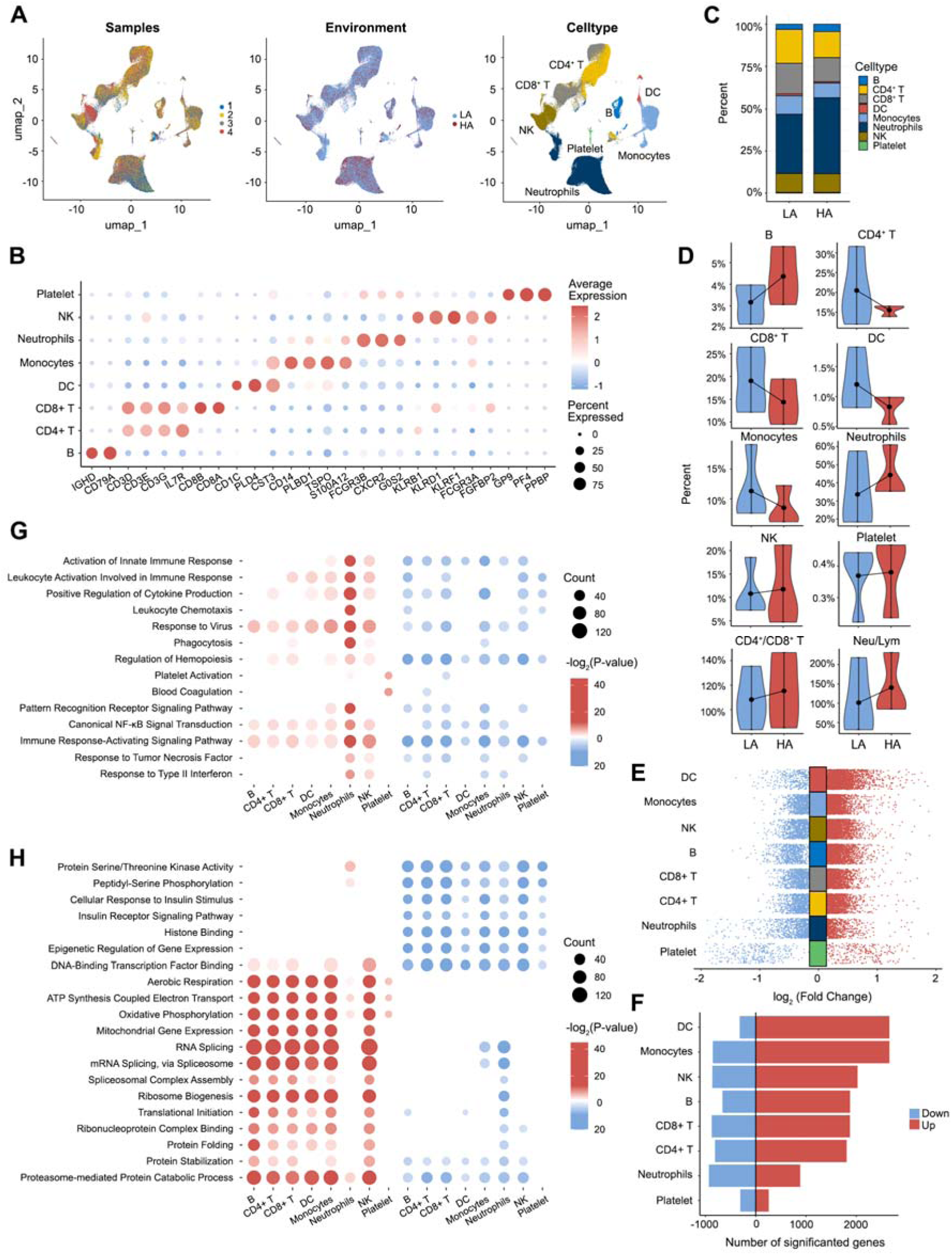
Comprehensive immune-metabolic profiling in response to LTHAE. Schematic overview of the experimental design and sample collection.

### 2. Single-cell RNA sequencing reveals LTHAE-induced immune landscape remodeling

To characterize the immune landscape associated with LTHAE, single-cell transcriptomic profiling of circulating immune cells from four subjects was conducted. After strict data quality control and standardized annotation process, a total of 125,844 high-quality cells were obtained, and their expression profiles covered 20,082 genes. Eight major cell types were further identified by clustering analysis and reference-based integration (Figure 2A). Cell type annotation was confirmed by classical marker gene expression profiles (Figure 2B)^21^.

**Figure 2.**
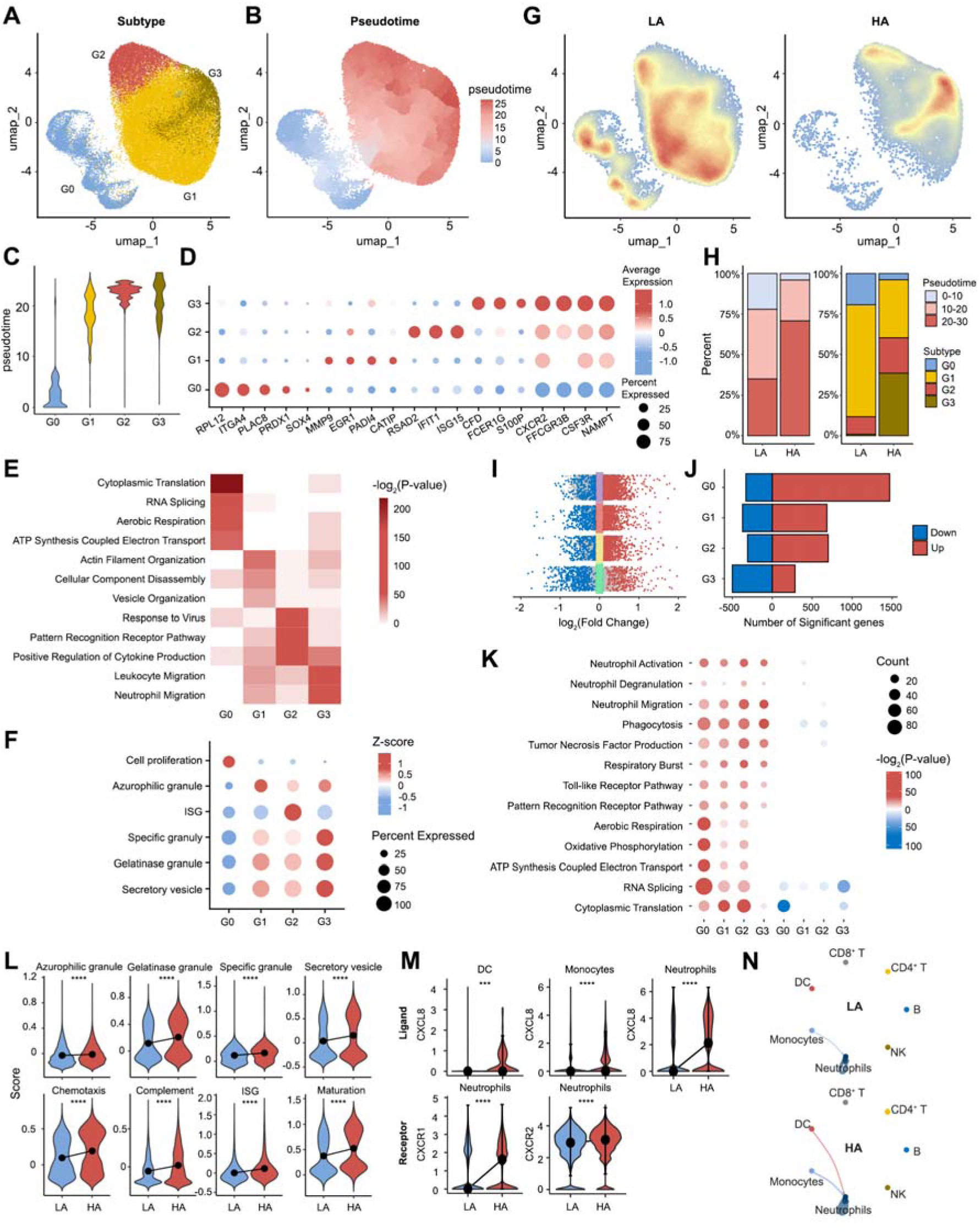
Single-cell RNA sequencing reveals LTHAE-induced immune landscape remodeling. A. UMAP analysis showing integrated patterns of sample attributes (left), environmental variables (middle), and eight immune cell types (right). B. Dot plots showing the percentages and average expressions of marker genes across eight immune cell types in the LA and HA groups. C-D. Bar plot(C) and violin plots(D) showing the changes in the relative proportions of immune cell types. E. Multi-group volcano plots displaying differentially expressed genes (DEGs) in eight immune cell types, with significantly upregulated (red) and downregulated (blue) genes indicated. F. Bar plot depicting the number of significant DEGs in each immune cell type. G-H. Functional enrichment analysis of DEGs reveals (G) immune-related pathways and (H) fundamental biological processes-related GO term profiles in eight immune cell populations. The bubble plots visualize significantly enriched GO terms, with color gradients representing -log2(P-values) for upregulated (red) and downregulated (blue) gene sets, and bubble size corresponds to the number of enriched genes.

Quantitative analysis at single-cell resolution revealed that the relative proportions of neutrophils, B lymphocytes, and natural killer (NK) cells in peripheral blood were increased following LTHAE, whereas those of CD4^+^ T cells, CD8^+^ T cells, dendritic cells (DCs), and monocytes were reduced (Figure 2C–D). Notably, both the neutrophil-to-lymphocyte ratio and the CD4^+^/CD8^+^ T cell ratio were found to increase, which was consistent with the immune remodeling pattern characterized by lymphocyte attenuation and myeloid cell expansion under high-altitude conditions that has been reported^22^. In addition, an increased proportion of platelets was observed, which may be associated with elevated thrombopoietin (TPO) levels and enhanced megakaryocyte proliferation induced by the high-altitude environment^23^. Differential gene expression profiling revealed that LTHAE led to upregulation of gene expression in lymphocytes (including NK, B, and T cells), monocytes, and DCs, suggesting that these cell subsets were in a state of transcriptional activation (Figure 2E–F). In contrast, gene expression changes in neutrophils and platelets were predominantly downregulated. Collectively, these findings indicated that both the composition of immune cell subsets and their transcriptomic features were substantially reshaped following LTHAE conditions. Enrichment analysis also showed that LTHAE led to functional remodeling of the immune system and changes in the basal metabolic process of cells. At the immune level, the immune function of B cells, CD4^+^ T cells, CD8^+^ T cells and monocytes showed an overall inhibition trend, while the activity of neutrophils was significantly enhanced, and the immune function of NK cells and DCs was partially activated (Figure 2G). These data suggest that hypoxia at high-altitude may preferentially activate the innate immune system and inhibit the adaptive immune response. At the molecular regulatory level, the observed alterations exhibited remarkable consistency across different immune cell types (Figure 2H). The insulin signaling pathway (involved in energy storage regulation), protein phosphorylation modifications (regulating enzyme activity), and epigenetic processes (controlling gene expression) were universally downregulated across multiple immune cell populations, suggesting that the body prioritizes survival under hypoxic conditions by suppressing non-essential metabolic activities. In contrast, pathways related to oxidative phosphorylation and ATP synthesis—key components of cellular energy metabolism—were significantly upregulated in all immune cells, indicating a compensatory enhancement of aerobic oxidation efficiency to maintain energy homeostasis. Additionally, processes such as RNA splicing, protein translation, folding, stabilization, and degradation were activated in six immune cell types, whereas these functions were reduced in neutrophils, suggesting that distinct immune subsets employ divergent strategies to cope with metabolic stress under high-altitude hypoxia.

### 3. A Multi-subset scRNA-seq analysis of neutrophil/NK/DC activation and monocyte/lymphocyte suppression

#### Cellular and functional heterogeneity in the neutrophil under LTHAE

As the highest proportion of immune cells in leukocytes, neutrophils play a key role in the maintenance of host defense and immune homeostasis under environmental stress^24^. Based on the cluster analysis and pseudo-time analysis, neutrophils were divided into four differentiation stages: G0 precursor cells at the beginning of differentiation, early mature cells at the intermediate transition stage of G1, and a G2/G3 mature functional subset at the end of differentiation (Figure 3A-B). G0 subpopulation specifically highly expressed precursor cell marker genes (RPL12, ITGA4, PLAC8), and their transcriptome characteristics were significantly enriched in basic metabolic pathways such as cytoplasmic translation, RNA splicing and aerobic respiration, accompanied by strong proliferation activity. With the progress of differentiation, G1 began to express maturation marker genes (MMP9, EGR1, PADI4, etc.), and its functional characteristics changed to actin dynamic remodeling and vesicle trafficking, accompanied by a large number of azurophilic granule synthesis. At the terminal differentiation stage, G2 subset was involved in pattern recognition and cytokine secretion through significant activation of interferon-stimulated genes (ISG). G3 played an important role in cell migration and pathogen clearance through the coordination of gelatinase granule, specific granule and secretory vesicle (Figure 3C-F).

**Figure 3.**
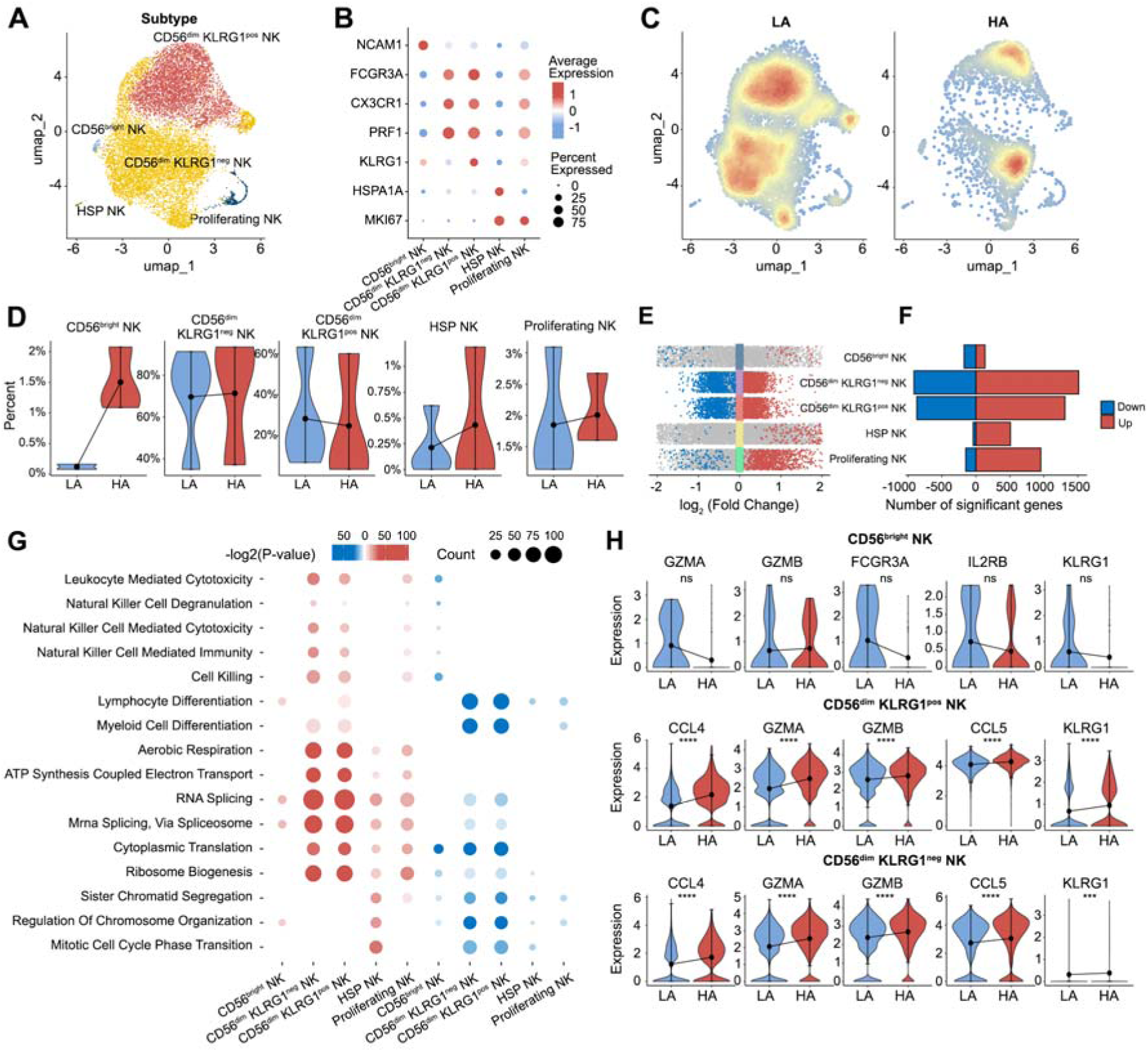
Cellular and functional heterogeneity in the neutrophil under LTHAE. A. UMAP plot of the neutrophil subtypes. B. Pseudotime analysis of neutrophil subtypes using Monocle 3. C. Violin plots showing the pseudotemporal distribution of neutrophil subtypes. D. Dot plots depicting the percentages and average expressions of marker genes in neutrophil subtypes. E. Gene ontology (GO) analysis of DEGs for each of the neutrophil subtypes. F. Bubble plot showing neutrophil-related signature scores calculated using the AddModuleScore algorithm. G. Cell density plots of neutrophils in LA and HA groups, with color intensity proportional to local cell density. H. Bar plots showing frequencies of neutrophil subtypes (left) and pseudotemporal distribution (right) in the LA and HA groups. I. Multi-group volcano plots displaying DEGs across neutrophil subtypes in LA and HA groups, with significantly upregulated (red) and downregulated (blue) genes highlighted. J. Bar plot depicting the number of DEGs in each neutrophil subtype. K. Functional enrichment analysis of DEG between LA and HA groups across neutrophil subtypes. The bubble plots visualize significantly enriched GO terms, with color gradients representing -log2(P-values) for upregulated (red) and downregulated (blue) gene sets, and bubble size corresponds to the number of enriched genes. L. Violin plots showing the changes in the neutrophil-specific signature scores in LA and HA groups. M. Expression distribution of CXCL signaling genes in the LA and HA groups, including the ligand CXCL8 and its receptors CXCR1 and CXCR2. N. The inferred CXCL signaling. Edge width represents the communication probability in each cell group.

LTHAE significantly changed the distribution of neutrophil subsets and functional status. Compared with the plain environment, the proportion of G0-G1 subsets in peripheral blood decreased and the proportions of G2–G3 mature subsets increased after plateau adaptation, while the distribution of pseudotime series scores shifted to higher values, suggesting that LTHAE may promote the differentiation of peripheral blood neutrophils to a more mature stage (Figure 3G–H). The differentially expressed genes (DEGs) in the G0–G2 subgroups were predominantly upregulated, whereas those in the G3 subgroup were mainly downregulated (Figure 3I–J). Enrichment analysis revealed that genes associated with immune functions—such as immune activation, phagocytosis, cytokine production, chemotaxis, respiratory burst, and degranulation—as well as signaling pathways including pattern recognition receptor and Toll-like receptor signaling, were significantly enriched across all G0–G3 neutrophil subsets (Figure 3K). Furthermore, functional gene set scoring indicated elevated expression of neutrophil granule-related genes, particularly those associated with secondary and tertiary granules. Functions linked to bactericidal and proinflammatory activities, including chemotaxis, ISG response, complement activation, and vesicle secretion, were also upregulated in neutrophils (Figure 3L). These findings suggested that LTHAE promotes neutrophil maturation and enhances their overall immune functionality.

Notably, although the immune functions of the G0–G3 subsets are all activated, there are differences in the molecular regulatory mechanisms. Enhanced oxidative phosphorylation and

RNA splicing were observed in the G0-G2 subset, whereas the G3 subset exhibited inhibited translation and reduced RNA splicing, reflecting the different adaptation strategies of cells at different differentiation stages to the plateau hypoxia environment (Figure 3K).

As a principal neutrophil chemotactic factor, CXCL8 is primarily secreted by monocytes and neutrophils, and it regulates neutrophil chemotactic migration and effector functions by binding to the CXCR1/2 receptors^25^. CellChat analysis revealed that the expression of CXCR1/2 on neutrophils was significantly upregulated following LTHAE. Meanwhile, the expression of CXCL8 in monocytes, neutrophils, and dendritic cells was synchronously increased (Figure 3M). The signaling communication intensity of the CXCL8–CXCR1/2 pathway was enhanced (Figure 3N), suggesting a potential role in the abnormal activation of neutrophils under hypoxic conditions.

#### Cellular and functional heterogeneity in the NK cell under LTHAE

After LTHAE, NK cells were identified as another cell population with increased proportion and enhanced immune activity. NK cells were classically defined as CD3□CD56□ lymphocytes and typically divided into two major subsets based on CD56 expression levels: CD56^bright^ (immature subset) and CD56^dim^ (terminally mature subset). Among these, the CD56^dim^ subset accounted for over 90% of peripheral NK cells. The cytotoxic activity, along with the expression level of the inhibitory receptor KLRG1, increases with cellular maturation. Based on their functional characteristics, NK cells were further classified into five subclusters (Figure 4A–B), including: CD56^bright^ NK cells (NCAM1□), CD56^dim^ KLRG1□ NK cells (FCGR3A□ CX3CR1□ PRF1□ KLRG1□), CD56^dim^ KLRG1□ NK cells (FCGR3A□ CX3CR1□ PRF1□ KLRG1□), Heat shock protein-associated NK cells (HSP NK; HSPA1A□ MKI67□), and proliferating NK cells (HSPA1A□ MKI67□). Consistent with previous studies, CD56^dim^ NK cells have been identified as the predominant population in peripheral blood^26^.

**Figure 4.**
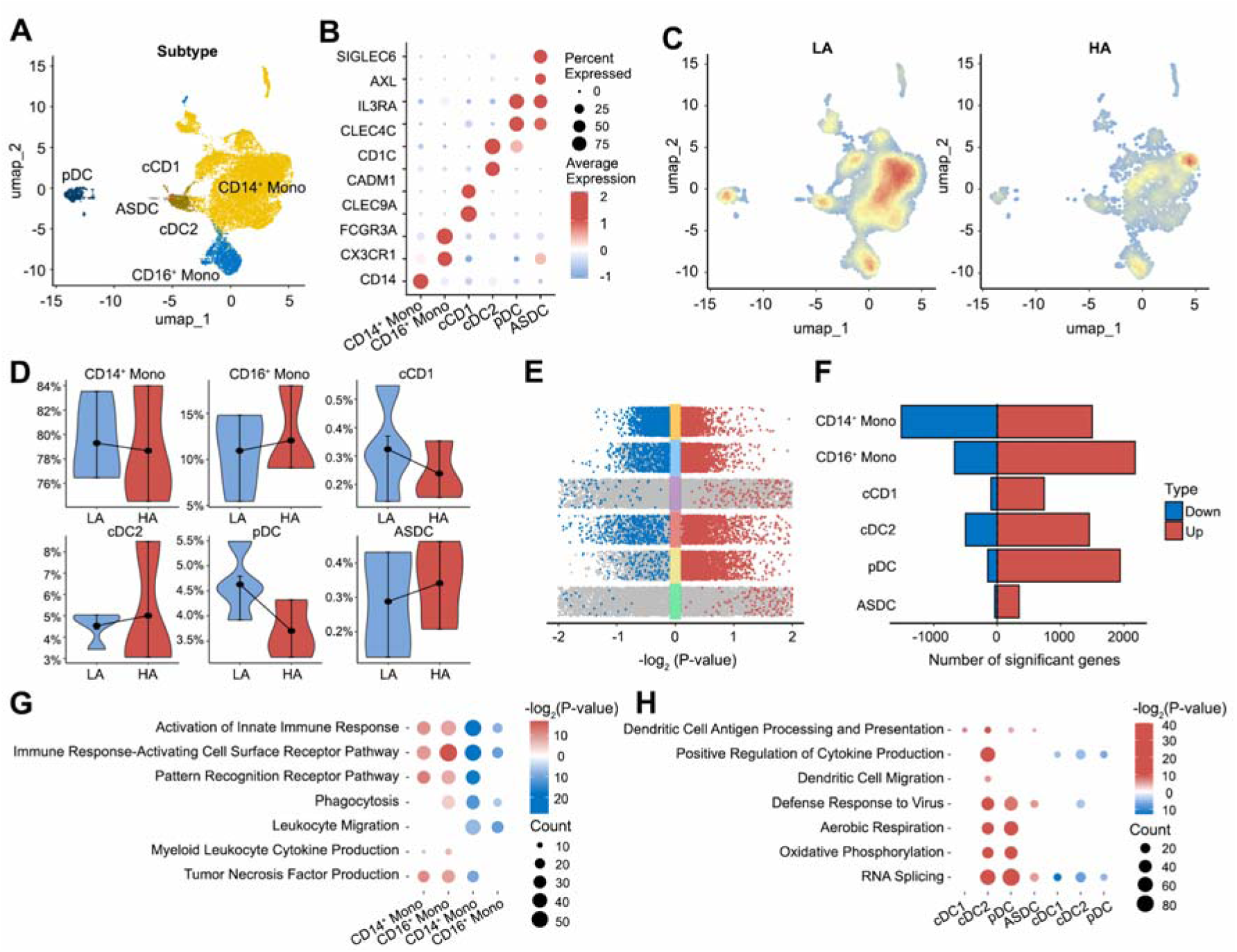
Cellular and functional heterogeneity in the NK cell under LTHAE. A. UMAP plot of the NK subtypes. B. Dot plots depicting the percentages and average expressions of marker genes in NK subtypes. C. Cell density plots of NK in LA and HA groups, with color intensity proportional to local cell density. D. Violin plots showing the changes in the relative proportions of NK subtypes in LA and HA groups. E. Multi-group volcano plots displaying DEGs across NK subtypes in LA and HA groups, with significantly upregulated (red) and downregulated (blue) genes indicated. F. Bar plot depicting the number of DEGs in each NK subtype in the LA and HA groups. G. Functional enrichment analysis of DEG between LA and HA groups across NK subtypes. The bubble plots visualize significantly enriched GO terms, with color gradients representing -log2(P-values) for upregulated (red) and downregulated (blue) gene sets, and bubble size corresponds to the number of enriched genes. H. Violin plots showing the immune-related genes in NK subtypes in LA and HA groups.

Compared to the LA group, LTHAE led to an increase in the proportions of CD56^bright^ NK cells and CD56^dim^ KLRG1□ NK cells (activated subsets), while the proportion of CD56^dim^ KLRG1□ NK cells (suppressive/terminally differentiated subset) decreased (Figure 4C–D). DEGs in the HSP NK, Proliferating NK, and CD56^dim^ NK subsets were predominantly upregulated, whereas DEGs in CD56^bright^ NK cells were mainly downregulated (Figure 4E–F). Enrichment analysis showed that immune-related genes involved in cytotoxicity, degranulation, and cell killing were significantly upregulated in both CD56^dim^ KLRG1□ and CD56^dim^ KLRG1□ NK subsets, whereas immune functional genes such as GZMA, FCGR3A, and IL2RB were broadly downregulated in the CD56^bright^ NK subset (Figure 4G-H).

Moreover, oxidative phosphorylation, RNA splicing, and protein translation were suppressed in the CD56^bright^ NK subset, while these processes remained active in other subsets. These findings suggested that LTHAE may reshape the immunological profile of NK cells by suppressing the immune and fundamental cellular functions of the CD56^bright^ NK subset and enhancing the cytotoxic activity of the CD56^dim^ NK subset.

#### Cellular and functional heterogeneity in the monocytes and DCs under LTHAE

Monocytes were clearly classified into two subsets based on the differential expression of CD14 and CD16 surface markers: classical monocytes (CD14□CD16□) and non-classical monocytes (CD14□CD16□) (Figure 5A–B). LTHAE reshaped both the subset distribution of monocytes. Compared with the LA group, the proportion of non-classical monocytes increased in the HA group (Figure 5C–D). Differential gene expression analysis (Figure 5E–F) and enrichment analysis (Figure 5G) revealed that genes related to phagocytosis and chemotactic migration were significantly downregulated in classical monocytes. In contrast, DEGs in non-classical monocytes were predominantly upregulated, accompanied by activation of the TNF-α signaling pathway. These findings indicated that LTHAE reshapes the immunological profile of monocytes by suppressing the activity of classical monocytes while enhancing the responsiveness of non-classical monocytes.

**Figure 5.**
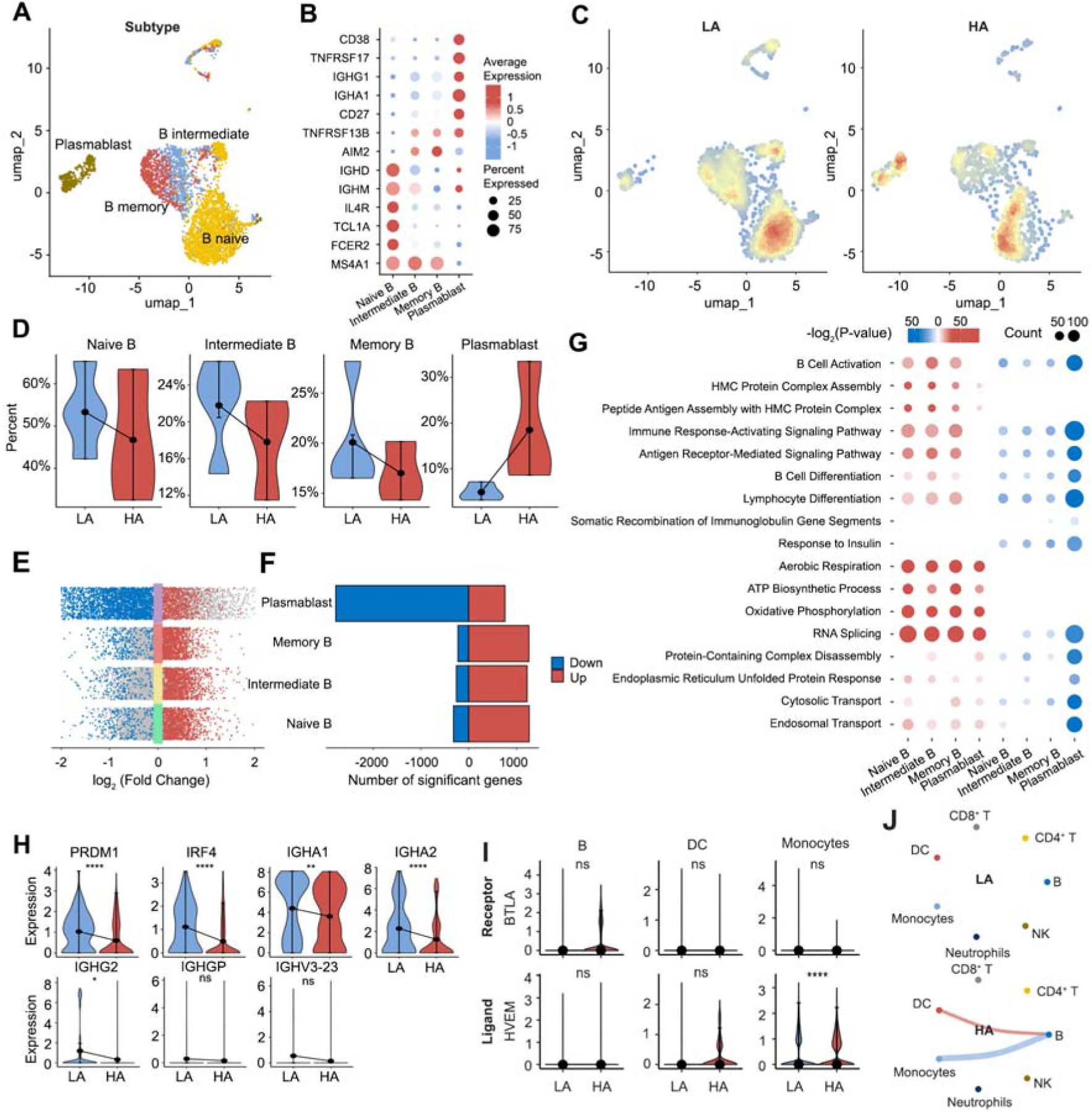
Cellular and functional heterogeneity in the monocyte and DCs under LTHAE. A. UMAP plot of the monocyte and DC subtypes. B. Dot plots depicting the percentages and average expressions of marker genes in monocyte and DC subtypes. C. Cell density plots of monocyte and DC in LA and HA groups, with color intensity proportional to local cell density. D. Volin plots showing the changes in the relative proportions of monocyte and DC subtypes in LA and HA groups. E. Multi-group volcano plots displaying DEGs across monocyte and DC subtypes, with significantly upregulated (red) and downregulated (blue) genes indicated. F. Bar plot depicting the number of DEGs in each monocyte and DC subtype. G. Functional enrichment analysis of DEGs between LA and HA groups across monocyte subtypes. The bubble plots visualize significantly enriched GO terms, with color gradients representing -log2(P-values) for upregulated (red) and downregulated (blue) gene sets, and bubble size corresponds to the number of enriched genes. H. Functional enrichment analysis of DEGs between LA and HA groups across DC subtypes.

DCs were classified into four functional subsets: plasmacytoid DCs (pDC; LILRA4□ IRF7□), AXL□ Siglec-6□ DCs (ASDC; AXL□ SIGLEC6□), classical type 1 DCs (cDC1; CD1C□ CLEC9A□), and classical type 2 DCs (cDC2; CD1C□ CLEC10A□) (Figure 5A–B).

Following LTHAE, the proportions of cDC2 and ASDC increased, while those of cDC1 and pDC decreased (Figure 5C–D). Differential gene expression analysis revealed that immune-related genes across all DC subsets were predominantly upregulated (Figure 5E–F). Enrichment analysis showed that the cDC2 subset exhibited enhanced cytokine secretion and chemokine receptor–mediated migration capacity, whereas the pDC subset displayed significant activation of antiviral response pathways. In addition, both subsets showed active oxidative phosphorylation and RNA splicing processes (Figure 5H). These findings indicated that the high-altitude environment enhances the innate immune responsiveness of both cDC2 and pDC subsets.

#### Cellular and functional heterogeneity in the B cell under LTHAE

As a crucial component of the adaptive immune response, B cells significantly influence the effectiveness of antibody-mediated immunity through their activation and differentiation status. In this study, B cells were classified into four distinct subsets (Figure 6A–B): naïve B cells (FCER2□ TCL1A□ IL4R□), intermediate B cells (IGHM□ AIM2□), memory B cells (IGHM□ AIM2□), and plasmablasts (CD38□ TNFRSF17□ IGHG1□). LTHAE led to an increase in the proportion of plasmablasts while decreasing the frequencies of naïve, intermediate and memory B cells (Figure 6C–D). These findings indicated that the high-altitude environment reshaped the developmental trajectory of B cells, driving their differentiation toward a plasma cell fate.

**Figure 6.**
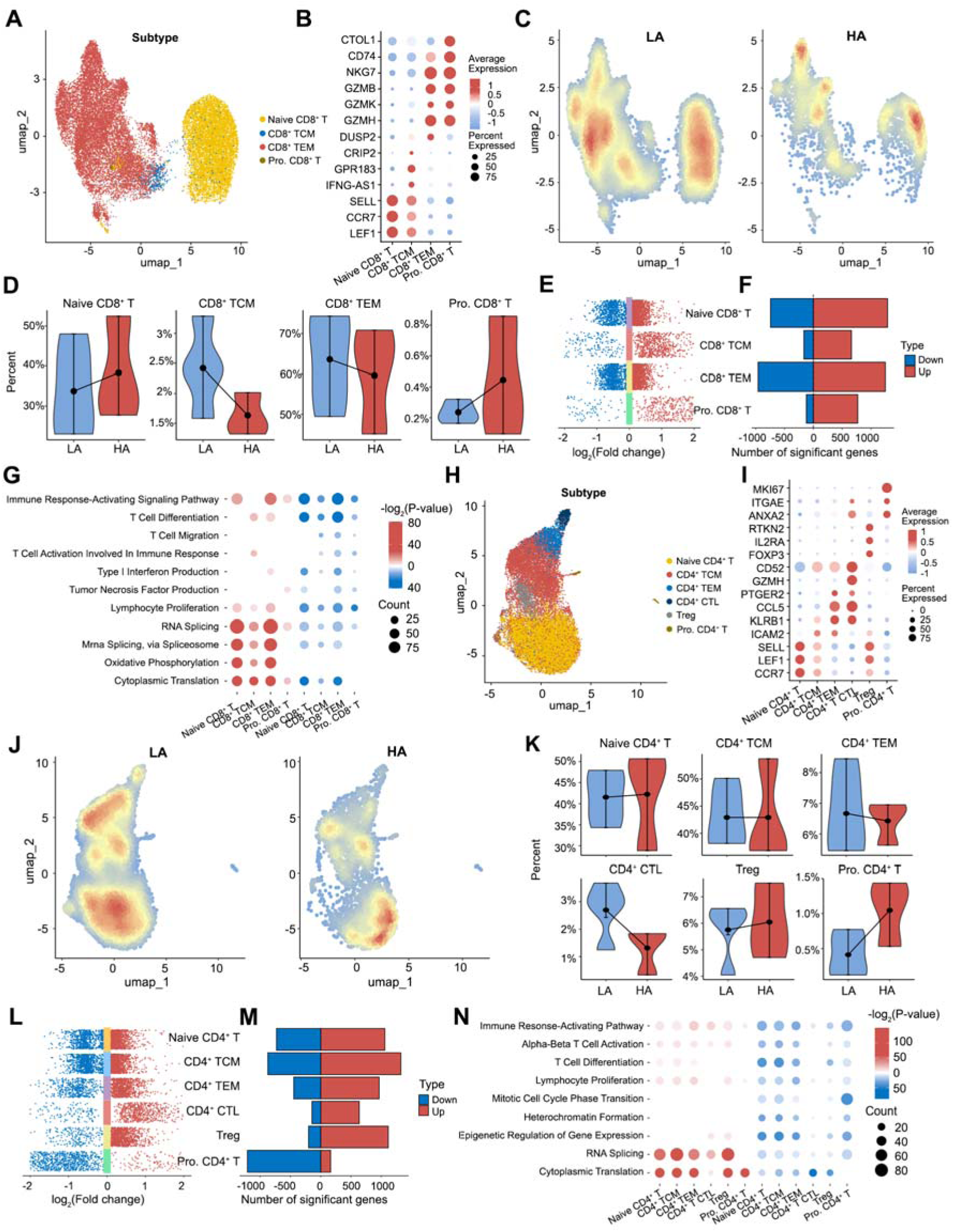
Cellular and functional heterogeneity in the B cell under LTHAE. A. UMAP plot of the B cell subtypes. B. Dot plots depicting the percentages and average expressions of marker genes in B cell subtypes. C. Cell density plots of B cell in LA and HA groups, with color intensity proportional to local cell density. D. Violin plots showing the changes in the relative proportions of B cell subtypes between LA and HA groups. E. Multi-group volcano plots displaying DEGs across B cell subtypes in LA and HA groups, with significantly upregulated (red) and downregulated (blue) genes indicated. F. Bar plot depicting the number of DEGs in each B cell subtype between LA and HA groups. G. Functional enrichment analysis of DEG between LA and HA groups across B cell subtypes. The bubble plots visualize significantly enriched GO terms, with color gradients representing -log2(P-values) for upregulated (red) and downregulated (blue) gene sets, and bubble size corresponds to the number of enriched genes. H. Violin plots showing the immune-related genes in plasmablast in LA and HA groups. I. Violin plots showing BTLA signaling genes in LA and HA groups. J. The inferred BTLA signaling. Edge width represents the communication probability in each cell group.

Differential gene expression analysis across the B cell subsets revealed that DEGs in naïve, intermediate and memory B cells were predominantly upregulated (Figure 6E–F), with functional enrichment in pathways related to B cell activation, antigen presentation, and pro-inflammatory signaling (Figure 6G), suggesting enhanced B cell differentiation and development. In contrast, DEGs in plasmablasts were mainly downregulated. Notably, the core molecular network regulating antibody secretion in plasmablasts was broadly suppressed, indicating a marked reduction in the functional activity of plasmablasts. Consistently, critical transcription factors required for plasma cell differentiation, such as PRDM1 and its co-factor IRF4, were significantly downregulated in plasmablasts, along with reduced expression of immunoglobulin genes (IGHA1, IGHA2, IGHG2) (Figure 6H). Additionally, genes involved in protein localization, complex assembly, and vesicle transport were also decreased (Figure 6G), indicating a comprehensive suppression spanning from transcriptional regulation, antibody synthesis, to secretory function in plasmablasts. These findings suggested that LTHAE might suppress plasmablast maturation and antibody production, specifically impairing humoral immune responses. At the same time, there appears to be a compensatory drive toward plasmablast differentiation (Figure 6D), possibly as a response to heightened immune challenges under hypoxic conditions.

As a key immunomodulatory ligand, HVEM is predominantly secreted by dendritic cells and monocytes. It suppresses cytokine production and costimulatory molecule expression in B cells by binding to the BTLA receptor ^27^. CellChat analysis showed that following LTHAE, the expression level of the BTLA receptor on B cells was upregulated, suggesting an increased sensitivity of B cells to BTLA signaling (Figure 6I). At the same time, the expression of HVEM ligands secreted by dendritic cells and monocytes was also elevated (Figure 6I). The coordinated upregulation of both ligand and receptor led to a significant enhancement in the communication intensity of the BTLA signaling pathway (Figure 6J), which may be associated with the suppression of B cell antibody production that has been observed under LTHAE.

#### Cellular and functional heterogeneity in the T cell under LTHAE

As another central component of adaptive immunity, T lymphocytes play essential roles in pathogen clearance and tumor immune surveillance. Based on classical surface markers, CD8□ T cells were classified into four functional subsets: naïve CD8□ T cells (CCR7□ TCF7□), proliferating CD8□ T cells (COTL1□ CAPG□), central memory CD8□ T cells (CD8□ TCM; GPR183□ CRIP2□), and effector memory CD8□ T cells (CD8□ TEM; DUSP□ GZMH□) (Supp. Fig. 1A–B). The CD8□ T cell population was predominantly composed of the naïve and effector memory subsets, consistent with previous studies ^28^.

LTHAE led to an imbalance in CD8□ T cell subsets (Supp. Fig. 1C–D), characterized by a decrease in overall frequency in peripheral blood (Figure 2C–D), depletion of memory subsets, and expansion of naïve and proliferating subpopulations. Differential gene expression and pathway enrichment analyses revealed that the differentiation and proliferation of the naïve subset were suppressed, while immune-related processes such as cell migration and cytokine secretion in the TEM subset were significantly impaired (Supp. Fig. 1E–G). These findings suggest that LTHAE may suppress the differentiation and/or functional capacity of various CD8□ T cell subsets.

CD4□ T cells were classified into six functional subsets: naïve CD4□ T cells (GPR18□ CD55□), proliferating CD4□ T cells (MKI67□ ITGAE□), central memory CD4□ T cells (CD4□ TCM; ICAM2□ GZMK□), effector memory CD4□ T cells (CD4□ TEM; ICAM2□ GZMK□), cytotoxic CD4□ T cells (CD4□ CTL; GZMH□ GZMA□), and regulatory T cells (Treg; FOXP3□ IL2RA□) (Supp. Fig. 1H–I). Following LTHAE, the overall proportion of CD4□ T cells in peripheral blood decreased (Figure 2C, D), with a notable reduction in CD4□ CTL and slight increases in both Treg and Proliferating CD4□ T cells (Supp. Fig. 1J– K). Differential gene expression and enrichment analyses revealed significant downregulation of processes related to T cell differentiation in the naïve subset, as well as impaired migration, immune activation and cytokine production in the TCM and TEM subsets (Supp. Fig. 1L–N).

These results suggest that LTHAE may suppress the differentiation potential and overall immune responsiveness of CD4□ T cells.

### 4. Untargeted metabolomics profiling reveals LTHAE-induced plasma metabolic landscape remodeling

To investigate the changes in plasma metabolites induced by LTHAE, untargeted metabolomics analysis of samples from the two groups was performed using ultra-performance liquid chromatography-electrospray ionization-quadrupole-orbitrap-mass spectrometry (UPLC-ESI-Q-Orbitrap-MS) (Supplementary Table 3), with a total of 2890 metabolites annotated. Principal component analysis (PCA) and orthogonal partial least squares discriminant analysis (OPLS-DA) revealed that the metabolite profiles of the two groups were significantly separated (Figure 7A-C), indicating that the body’s metabolic processes had changed significantly after LTHAE. By variable importance projection (VIP>1) and differential metabolomic analysis (P<0.05), 863 up-regulated metabolites and 143 down-regulated metabolites were identified (Figure 7D). KEGG pathway enrichment analysis showed that pathways such as caffeine metabolism, steroid hormone synthesis, unsaturated fatty acid metabolism, and amino acid metabolism were significantly activated (Figure 7E). Specifically, plasma levels of caffeine metabolites, steroid hormones, and polyunsaturated fatty acids were increased, while amino acid metabolism exhibited heterogeneity (with up-regulated histidine and tryptophan and down-regulated tyrosine and isoleucine) (Figure 7F).

**Figure 7.**
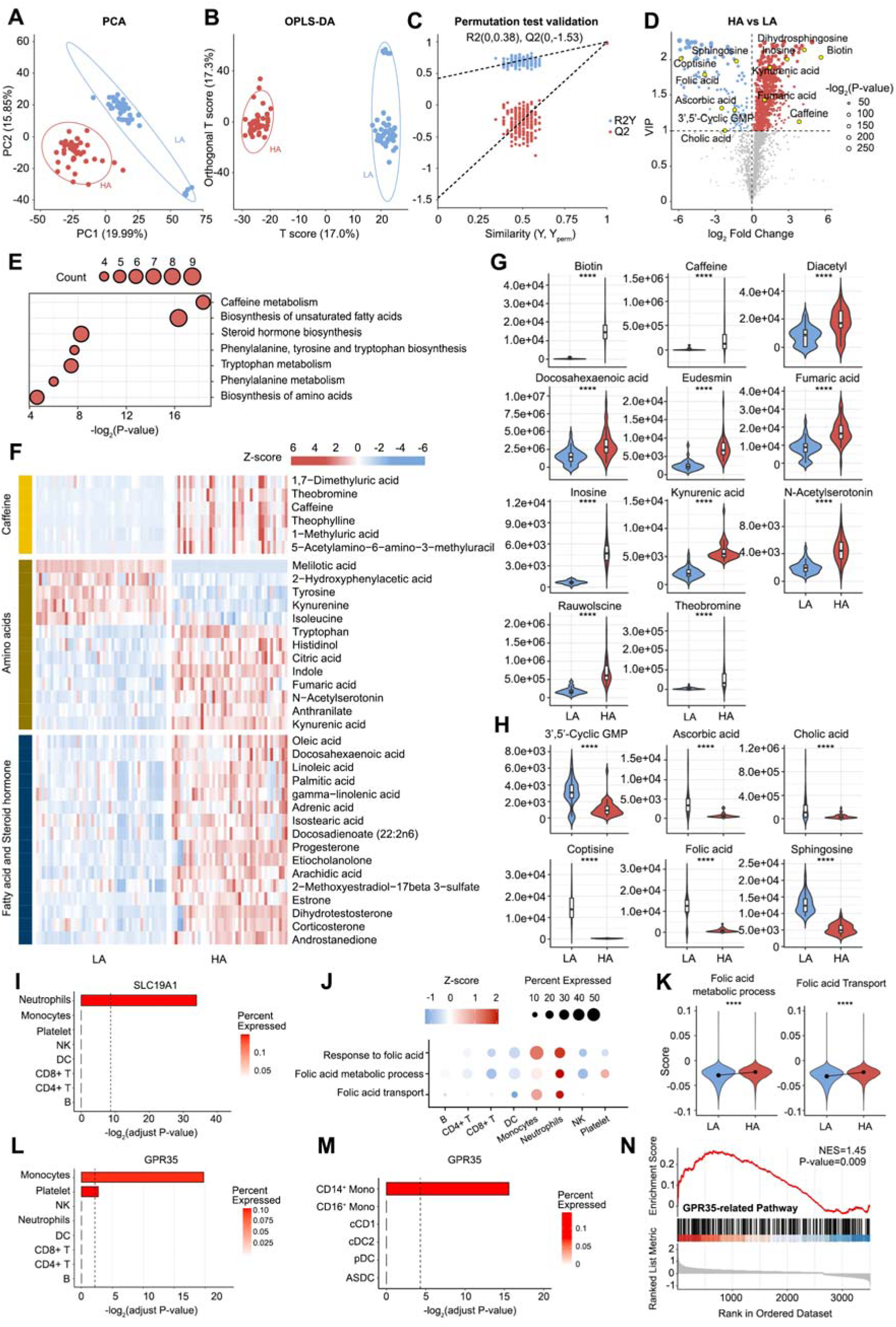
Untargeted metabolomics profiling reveals LTHAE-induced plasma metabolic landscape remodeling. A. PCA plot showing the LA (red dots) and HA (blue dots) samples were well separated. B-C. OPLS-DA analysis (B) and displacement test (C) of LA and HA samples. D. Volcano plot illustrating significantly differential metabolites (VIP>1 & P-value<0.05) between LA and HA groups. Metabolites with no statistical significance change (gray dots), significantly upregulated (red) and downregulated (blue) are indicated. E. Bubble plot of KEGG pathway enrichment analysis derived from significantly differential metabolites. Pathways are ranked by -log2(P-value), with bubble size representing the number of enriched metabolites indicating the significance level. F. Heatmap of significantly differential metabolites associated with caffeine metabolism, amino acids metabolism, fatty acids metabolism and steroid hormones biosynthesis. G-H. Violin plots showing the immunomodulatory metabolites that were significantly upregulated (G) and downregulated (H) between LA and HA groups, with statistical significance determined by two-tailed unpaired t-tests. I. Bar plots showing upregulated SLC19A1 expression across eight immune cell types, displaying mean expression percentages and adjusted p-values. J. Bubble plot showing folic acid-related signature scores. The color intensity represents the relative expression levels (Z-scores) of signature scores, while the bubble size indicates the percentage of expressing cells for each signature within cell types. K. Violin plots showing the changes in the folic acid-related signature scores of neutrophils between LA and HA groups. L-M. Bar plots showing upregulated GPR35 expression across eight immune cell types (L) and monocyte and DC subtypes (M), displaying mean expression percentages and adjusted p-values. N. GSEA reveals positive enrichment of the GPR35-related pathway in DEGs of monocytes (gene set derived from GSE15301).

### 5. LTHAE-induced plasma metabolic remodeling can influence the immunological functions

Emerging evidence have shown that changes in plasma metabolites can directly modulate immune cell function ^14^. To investigate potential impact of plasma metabolic remodeling on immunological outcomes, metabolites that met strict screening criteria were identified using the KEGG database (fold change >2 or <0.5, VIP > 1, P < 0.05), among which 57 were significantly up-regulated and 29 were significantly down-regulated (Supplementary Table 3). Among the up-regulated metabolites, 17 have been confirmed to have immune regulatory effects, including 11 anti-inflammatory metabolites (such as biotin, caffeine, fumarate and kynurenic acid) (Figure 7G). These have been demonstrated to regulate immune homeostasis by inhibiting the NF-κB/MAPK signaling pathway and reducing TNF-α/IL-6 proinflammatory factor expression^29–32^. Correspondingly, 3’,5’ -cyclic guanosine phosphate (cGMP) and folic acid, among the down-regulated metabolites (Figure 7H) (Supplementary Table 4), participate in the two-way regulation of inflammation through a complex immune regulatory network, and their reduction may weaken the proinflammatory signal ^33, 34^. These results suggest that LTHAE may influence immune homeostasis by modulating the balance between anti-inflammatory and pro-inflammatory metabolites.

As a core metabolite of nucleic acid synthesis and methylation modification, folic acid has been widely reported for its immunomodulatory function^35^. The analysis revealed that LTHAE significantly reduced plasma folic acid levels (Figure 7H), potentially impairing folic acid-dependent processes such as DNA repair and immune cell proliferation. However, neutrophils maintained a more active folic acid metabolic pathway than other immune cells by specifically upregulating the expression of the folic acid transporter SLC19A1 (Figure 7I-J). And the folic acid metabolic activity of neutrophils further increased from their own baseline before exposure (Figure 7K). This compensatory regulation might be closely related to the activation of neutrophils under hypoxic conditions.

The tryptophan metabolite kynurenic acid (KA) is a critical regulatory molecule in immune homeostasis and has been shown to suppress monocyte inflammatory responses by activating the G protein–coupled receptor GPR35^31^. Notably, plasma KA levels were significantly elevated under high-altitude conditions, by approximately 2.8-fold (Figure 7G), suggesting a potential role in modulating immune adaptation to hypoxia. Moreover, the upregulation of GPR35 expression was specifically observed in the classical monocyte subset (Figure 7L-M), indicating enhanced KA pathway signaling. Consistently, Gene set enrichment analysis (GSEA) revealed significant enrichment of genes associated with GPR35 downstream signaling pathways in classical monocytes under hypoxic conditions (Figure 7N). These findings suggested that hypoxia at high altitudes may lead to KA accumulation, which in turn specifically activates GPR35 on classical monocytes, ultimately mediating the suppression of their immune function.

### 6. Transcriptomic-metabolic integration analysis reveals LTHAE enhances aerobic oxidation efficiency

Previous studies have shown that short-to-medium-term (3–16 days) high-altitude exposure activates glycolysis and the pentose phosphate pathway ^18, 36, 37^. To investigate the effects of LTHAE on energy metabolism, 1,229 consistently up-regulated genes and 457 consistently down-regulated genes were screened by integrating the DEGs of immune subsets including neutrophils, NK cells, monocytes, DCs, B cells and T cells (Figure 8A-B). Enrichment analysis results showed that metabolic processes such as oxidative phosphorylation, citrate cycle (TCA cycle), glycolysis and gluconeogenesis were active, while lysine degradation and insulin resistance were inhibited (Figure 8C), suggesting that aerobic oxidation and energy metabolism were generally activated after LTHAE.

**Figure 8.**
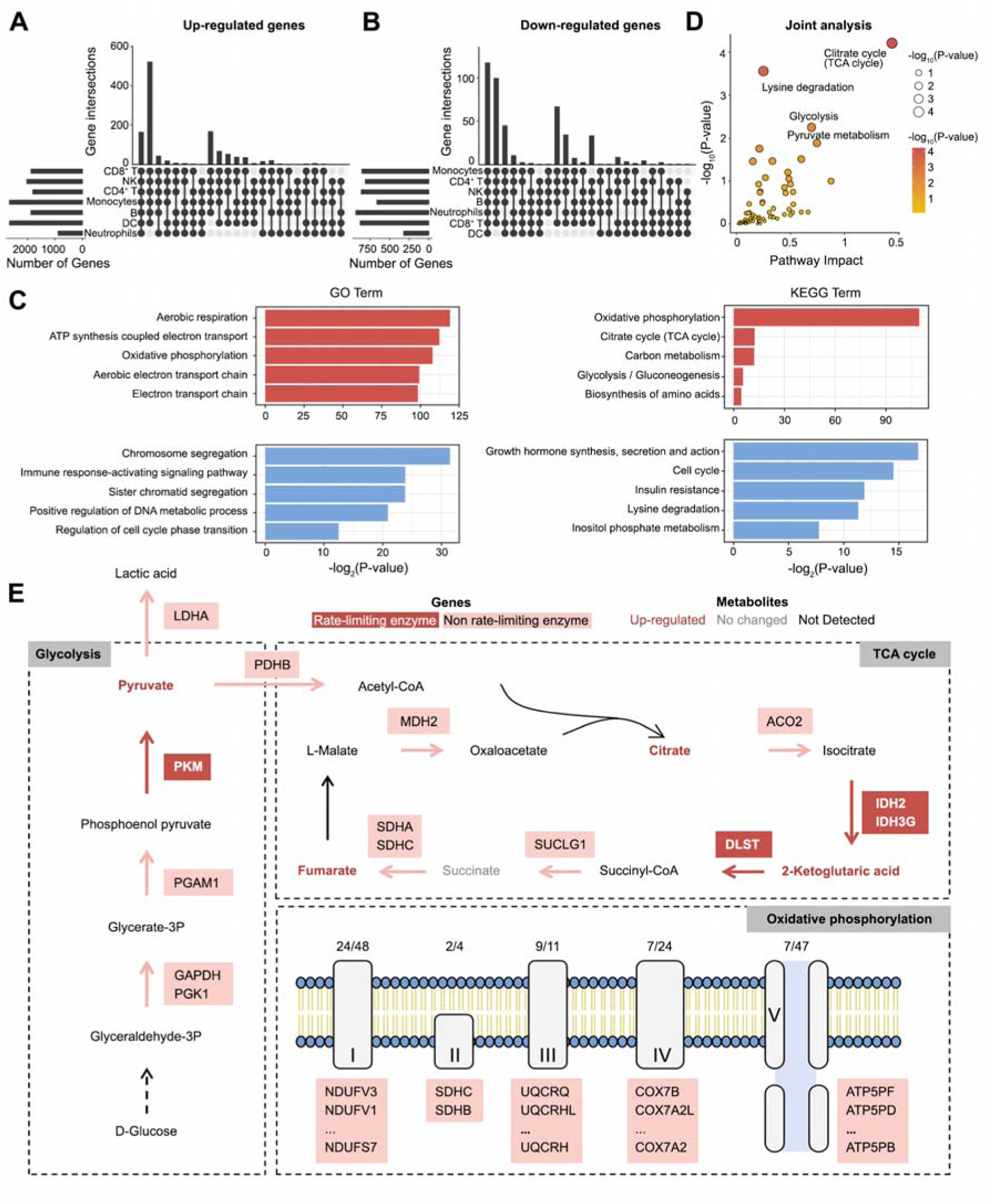
Transcriptomic-metabolic integration analysis reveals LTHAE enhances aerobic oxidation efficiency. A-B. The upset plot showing the intersection patterns of (A) upregulated and (B) downregulated DEGs among seven characterized immune cell populations. C. Bar plots displaying significantly enriched Gene Ontology terms and KEGG pathways among co-upregulated (red) and co-downregulated (blue) DEGs. D. Joint analysis of metabolomics data and coordinately changed single-cell transcriptomics. E. The metabolic pathway schematic depicts coordinated alterations in genes and metabolites across core energy metabolism pathways (glycolysis, TCA cycle, and oxidative phosphorylation). Upregulated genes are displayed with filled colors, where dark red denotes rate-limiting enzymes and light red represents non-rate-limiting enzymes. Metabolites are shown as unfilled shapes, color-coded as follows: red for significant elevation, gray for no change, and black for undetected.

To elucidate the mechanisms underlying the enhanced aerobic oxidative efficiency induced by LTHAE, we performed an integrated transcriptomic and metabolic analysis. The results showed that the multi-level enhancement of glycolysis-citric acid cycle-oxidative phosphorylation axis may be the core feature of metabolic adaptation after LTHAE (Figure 8D). The gene expression of pyruvate kinase (PKM), the rate-limiting enzyme of glycolysis, was up-regulated synchronously with pyruvate metabolites, drove the acceleration of glucose catabolism. Increased expression of pyruvate dehydrogenase (PDHB) promoted the conversion of pyruvate to acetyl-coa. The gene expressions of rate-limiting enzymes isocitrate dehydrogenase (IDH)2, IDH3G and α-ketoglutarate dehydrogenase (DLST) in TCA cycle were synergistically increased with the levels of intermediate metabolites such as citrate and α-ketoglutarate, indicating that the flux of TCA cycle was significantly increased. Extensive up-regulation of key complexes of mitochondrial oxidative phosphorylation further enhanced the efficiency of the electron transport chain and maximized ATP production per unit oxygen consumption. In addition, elevated lactate dehydrogenase (LDHA) expression suggested activation of an alternate pathway of the glycolytic-lactate cycle that may maintain metabolic resilience during fluctuations in oxygen supply (Figure 8E).

## Discussion

This study systematically characterized the dynamic immune cell regulation landscape and metabolic remodeling features in lowland populations following LTHAE, through integrated peripheral blood single-cell transcriptomic and plasma metabolomic analyses. The findings highlight potential complex interactions between immune and metabolic systems under hypoxic conditions, providing new clues into the molecular mechanisms underlying high-altitude adaptation.

Previous studies have revealed the staged characteristics of metabolic reprogramming under high-altitude hypoxia exposure: during the acute phase (6 hours), energy compensation is primarily driven by fatty acid β-oxidation^38^. In the short-to-medium term (3-16 days), mitochondrial function is suppressed to reduce ROS accumulation, while glycolysis and the pentose phosphate pathway are activated^18, 36, 37^. Notably, indigenous high-altitude populations, such as the Sherpas, have evolved unique genetic adaptations^17^. Their low expression of the PPARA gene suppresses fatty acid β-oxidation and activates the ω-oxidation pathway, converting fatty acids into succinate which is directly fed into the downstream TCA cycle. This mechanism avoids lipotoxicity while enhancing aerobic efficiency. However, the mechanisms underlying the establishment of metabolic homeostasis in long-term high-altitude adaptation, particularly the molecular basis for systemic optimization of metabolic networks in lowland populations, remain unclear. This study revealed that LTHAE (90 days) is characterized by systemic activation of the glycolysis-TCA cycle-oxidative phosphorylation axis. Single-cell transcriptomic analysis showed significant upregulation of genes encoding key metabolic enzymes, while plasma metabolomics detected synchronous accumulation of critical intermediates such as pyruvate and citrate. This transition from an emergency energy supply mode to a homeostatic optimization strategy suggests that hypoxic adaptation may be achieved through extensive remodeling of the metabolic network—a potential evolutionary response to sustained environmental stress. Moreover, both lowland individuals following LTHAE and indigenous high-altitude populations exhibit elevated LDHA expression, highlighting the conserved role of the glycolysis-lactate cycle in buffering oxygen availability during adaptation to hypoxia^17^.

A growing body of evidence suggests that acute high-altitude exposure rapidly activates neutrophil chemotaxis and macrophage phagocytic function via the HIF-1α signaling pathway, forming the first line of innate immune defense^39^. However, LTHAE may lead to functional suppression of T and B lymphocytes, potentially disrupting immune homeostasis and increasing susceptibility to infections^40^. Currently, there remains a lack of systematic understanding regarding the dynamic developmental trajectories of immune cell subsets and their functional regulation during long-term high-altitude adaptation. In this study, single-cell transcriptomic analysis was employed to characterize the remodeling of the peripheral immune system following LTHAE. Significant expansion and functional activation of myeloid cell populations were observed, in striking contrast to the proportional decline of lymphoid cells as well as specific monocyte-dendritic cell subsets. This regulatory mode is consistent with the acute high-altitude response reported by previous studies^19, 22^, but its molecular mechanism shows more complex metabolic-immune interaction characteristics in long-term adaptation.

Neutrophils are among the first cells of the innate immune system to be recruited to sites of infection and inflammation, playing a critical role in host defense against microbial pathogens^24^. Previous studies have shown that sustained hypoxia exerts profound effects on neutrophil function and survival, characterized by increased degranulation and prolonged lifespan^41^. LTHAE was associated with a significant increase in the proportion of peripheral blood neutrophils progressing toward terminal maturation stages (G1–G3), accompanied by widespread activation of granule synthesis, chemotactic migration, and antimicrobial effector functions. This functional profile may exert dual biological effects on the host. On one hand, the expansion of terminally mature subsets and enhanced antimicrobial activity may significantly improve early defense against pathogen infections. On the other hand, the extended survival and persistent degranulation of neutrophils may increase the risk of protease-mediated tissue damage^42^. An important finding of this study is the adaptive regulation of folic acid metabolism under high-altitude conditions. Plasma folic acid levels were markedly reduced following LTHAE, a result highly consistent with recent epidemiological studies conducted in Tibetan populations^43^. This suggests that systemic folic acid deficiency may represent a common metabolic feature of the hypoxic high-altitude environment. Although folic acid depletion may constrain immune cell proliferation and epigenetic regulation, neutrophils specifically upregulate SLC19A1-mediated folic acid uptake, thereby maintaining—and even enhancing—nucleic acid synthesis and methylation capacity under hypoxic stress^44, 45^. This adaptation likely contributes to the optimization of neutrophil function in response to chronic oxygen deprivation.

Classical and non-classical monocyte subsets perform distinct biological functions. Classical monocytes are characterized by their phagocytic capacity, chemotactic migration, and secretion of pro-inflammatory cytokines, and they serve as a “vanguard” in acute inflammation and tissue repair. In contrast, non-classical monocytes are generally considered to possess anti-inflammatory and immune surveillance functions, contributing to the resolution of inflammation through the production of anti-inflammatory cytokines and suppression of inflammatory signaling pathways^46^. Studies on short-term high-altitude exposure have shown that the proportion of classical monocytes increases rapidly during the acute phase, followed by a significant rise in intermediate monocytes on day 3^19^. These findings suggest that the body responds to acute inflammatory stress by rapidly mobilizing the phagocytic and chemotactic functions of classical monocytes. In contrast, this study revealed that LTHAE leads to a decrease in the proportion of classical monocytes and a marked increase in non-classical monocytes. Notably, specific upregulation of the GPR35 receptor on classical monocytes was observed alongside elevated plasma levels of kynurenine. This indicates that reprogramming of tryptophan metabolism may suppress the pro-inflammatory activity of classical monocytes via the kynurenine–GPR35 axis^31^. The dynamic shift from classical monocyte expansion to non-classical dominance suggests that chronic hypoxia may reduce the capacity of monocytes to infiltrate inflammatory sites, while promoting anti-inflammatory and immune surveillance functions.

Dendritic cells, as a critical link between innate and adaptive immunity, perform distinct roles in antigen presentation and immune regulation through their diverse subsets^47^. Among them, cDC1 cells activate CD8□ T cell-mediated anti-tumor immunity through cross-presentation, while cDC2 cells predominantly drive CD4□ T cell activation. pDCs participate in antiviral defense primarily through type I interferon responses, and ASDCs are potentially involved in immune tolerance through regulatory functions^48^. This study found that LTHAE led to a significant increase in the proportions of cDC2 and ASDC subsets, accompanied by a decrease in cDC1 and pDC populations. Enhanced chemokine receptor–mediated migration capacity and upregulated cytokine secretion were observed specifically in the cDC2 subset, while antiviral signaling pathways in pDCs were activated. These findings suggest a marked inclination toward innate immune activation in the host response under chronic hypoxia. This functional pattern, characterized by enhanced innate immunity, may represent a compensatory strategy adopted by the body under hypoxia.

Previous studies have shown that short-term high-altitude exposure (1–3 days) rapidly increases the frequency of total B cells in circulation, whereas prolonged hypoxia may lead to functional exhaustion of B cells, characterized by reduced proliferation capacity and impaired antibody secretion^19^. This study further elucidates the impact of long-term high-altitude hypoxia on B cell differentiation and function. A global suppression of antibody secretion–related genes in plasmablasts suggests compromised immune functionality, while their compensatory expansion may reflect an attempt by the host to enhance B cell differentiation and partially offset functional deficits^49^. Notably, upregulation of genes involved in B cell activation, antigen presentation, and pro-inflammatory signaling pathways in naïve and transitional B cell subsets indicates that the host may attempt to compensate for impaired antibody production by enhancing early immune recognition^50, 51^. This functional imbalance may exert dual effects on adaptive immunity under high-altitude conditions. On one hand, sustained suppression of plasmablast antibody secretion could delay or weaken humoral immune responses against extracellular pathogens, thereby increasing susceptibility to infections at high altitudes. On the other hand, the reduction in memory B cell frequency may impair the formation of long-term immunological memory, potentially diminishing vaccine efficacy or the responsiveness to repeated pathogen encounters^52^. Importantly, although the hyperactivation of naïve and transitional B cells may partially compensate for antibody deficiency, persistent activation of pro-inflammatory signaling pathways may disrupt immune tolerance, potentially increasing the risk of autoimmune disorders under chronic hypoxia^53^. Furthermore, the functional dissociation among B cell subsets—enhanced early activation but suppressed terminal differentiation—may compromise the coordination of immune responses, ultimately reducing the host’s overall defense capability against complex pathogens. As another core component of adaptive immunity, the functional suppression of both CD8□ and CD4□ T cells further support the notion of a “strategic contraction” of adaptive immunity under high-altitude hypoxia. The depletion of memory CD8□ T cell subsets and impaired effector functions may hinder the clearance of intracellular pathogens, while the widespread downregulation of activation-related genes in CD4□ T cells likely contribute to weakened coordination of immune responses^54^. The diminished effector function of T cells combined with defective B cell antibody production may create a “dual deficit,” leading to a substantial decline in the overall effectiveness of adaptive immune responses against complex pathogens in high-altitude environments^39^.

Although this study provides insights into the metabolic and immune characteristics associated with LTHAE, key limitations warrant mention. The relatively small sample size and exclusive focus on LTHAE individuals, without short-term exposure comparisons, constrain the ability to fully discern long-term adaptation mechanisms and assess population heterogeneity. Furthermore, while single-cell transcriptomics revealed transcriptional dynamics, the absence of functional validation (e.g., antibody secretion or cytotoxic activity assays) limits causal links to functional phenotypes. Critically, obtaining samples suitable for such functional assays under the challenging conditions of high-altitude field studies proved exceptionally difficult due to logistical constraints in sample collection, preservation, and immediate processing. Future studies should aim to expand cohort sizes, include control groups of individuals with short-term exposure, and incorporate functional assays to systematically dissect the interactions between metabolism and immunity. Further elucidation of these mechanisms will help uncover their pathological implications in high-altitude disorders, such as high-altitude pulmonary edema (HAPE) and chronic mountain sickness, thereby advancing our understanding of hypoxia-related disease progression and adaptation.

## Materials and Methods

### Sample collection and ethics statement

This study was conducted at two distinct geographical sites: Jiangjin, Chongqing (200 m above sea level) representing a plain area, and Lhasa, Tibet (3,650 m above sea level) as a typical moderate-altitude plateau region. Lhasa provides significant hypoxia with atmospheric oxygen levels at ∼60–65% of sea level, offering a safe model for studying human adaptation without high risk of acute mountain sickness.

A total of 46 healthy male Chinese volunteers residing in low-altitude regions (<300 m) with no history of chronic diseases were recruited. Baseline physiological assessments and fasting venous blood samples were collected from 45 participants at the plain site (200 m). After relocating to Lhasa (3,650 m), 40 participants completed follow-up evaluations following a 90-day acclimatization period.

To investigate immune homeostasis in LTHAE, a subgroup of 4 male volunteers was randomly selected from the original cohort of 46 healthy individuals. Fasting venous blood samples were obtained at the plain site from all 4 volunteers for peripheral blood single-cell RNA sequencing, with repeated sampling after 90 days of acclimatization in Lhasa (3,650 m); one participant was subsequently excluded during the high-altitude phase due to non-study-related factors. This study was approved by the Institute Research Medical Ethics Committee of the Army Medical University. A written informed consent was obtained from each participant.

### Metabolomic profiling and data processing

#### Sample collection and UHPLC-MS/MS analysis

Whole blood was collected in 4 mL heparin tubes, centrifuged (1300×g, 10 min), and plasma was aliquoted (0.2 mL/tube), flash-frozen in liquid nitrogen, and stored on dry ice. For analysis, thawed plasma (4 °C) was mixed with chilled methanol (1:3), incubated (-20 °C, 30 min), centrifuged (16,000×g, 20 min, 4 °C), and the supernatant was analyzed by subsequent LC-MS analysis.

Metabolomics profiling was analyzed using a UPLC-ESI-Q-Orbitrap-MS system (UHPLC, Shimadzu Nexera X2 LC-30AD, Shimadzu, Japan) coupled with Q-Exactive Plus (Thermo Scientific, San Jose, USA). UPLC-MS/MS analysis was performed on a Waters ACQUITY UPLC coupled to an TripleTOF® 6600 mass spectrometer (AB SCIEX) equipped with an ESI source. Chromatographic separation was performed on aWaters C18 column (2.1×100 mm, 1.6 µm).

Raw data were processed using MSDIAL software for peak alignment, retention time correction, and peak area extraction. Metabolite identification was performed through accurate mass matching (mass tolerance < 10 ppm) and MS/MS spectral matching (mass tolerance < 0.01 Da) against public databases (HMDB, MassBank, GNPS) and the BIOPROFILE-Database. For statistical analysis, ion peaks with >50% missing values within groups were excluded. The remaining data were normalized by total peak area separately for positive and negative ion modes, followed by integration of dual-mode features. Python software was employed for pattern recognition analysis after Unit Variance Scaling (UV) preprocessing.

#### Multivariate statistical analysis

Data were preprocessed using Pareto scaling and analyzed via principal component analysis (PCA), and partial least-square discriminant analysis (OPLS-DA). All the models evaluated were tested for over fitting with methods of permutation tests(n=200), with performance evaluated by R2X(cumulative), R2Y(cumulative), and Q2(cumulative) metrics. Differentially abundant metabolites were identified based on VIP scores (>1.0) from OPLS-DA and p-values (<0.05) from two-tailed Student’s t test. Fold change was calculated as the logarithm of the average mass response (area) ratio between two arbitrary classes.

#### KEGG enrichment analysis of differential abundant metabolites

The differentially abundant metabolites were performed KEGG pathway analysis using KEGG database (http://www.kegg.jp). KEGG enrichment analyses were carried out with the hypergoemetric test, and FDR correction for multiple testing was performed. Enriched KEGG pathways were nominally statistically significant at the p<0.05 level.

### Single-cell sequencing and data processing

#### Sample preparation of single-cell suspensions

Whole blood samples were collected in EDTA tubes and transported to the laboratory under cold chain conditions. For red blood cell lysis, 0.5 mL of whole blood was diluted with PBS (HyClone) to 10 mL, followed by the addition of 3 volumes of red blood cell lysis buffer (RCLB, Singleron) to the cell suspension. After incubation at room temperature for 10-12 minutes, the mixture was centrifuged at 450×g for 10 minutes. The supernatant was discarded, and the cell pellet was resuspended in 1 mL PBS. After an additional PBS washing step, the suspension was centrifuged at 300×g for 5 minutes. The supernatant was removed, and the cell pellet was resuspended in an appropriate volume of PBS. Cell viability and concentration were determined using the Countstar® Rigel fluorescence cell counter with AO/PI staining.

#### Sequencing library construction

Single-cell suspensions (2×10^5^ cells/mL) with PBS were loaded onto microwell chip using the Singleron Matrix® Single Cell Processing System. Barcoding Beads are subsequently collected from the microwell chip, followed by reverse transcription of the mRNA captured by the Barcoding Beads and to obtain cDNA, and PCR amplification. The amplified cDNA is then fragmented and ligated with sequencing adapters. The scRNA-seq libraries were constructed according to the protocol of the GEXSCOPE® Single Cell RNA Library Kits (Singleron) ^55^. Individual libraries were diluted to 4 nM, pooled, and sequenced on Illumina novaseq 6000 with 150 bp paired end reads.

#### Primary analysis of raw read data

Raw reads from scRNA-seq were processed to generate gene expression matrixes using CeleScope (https://github.com/singleron-RD/CeleScope) v1.9.0 pipeline. Briefly, raw reads were first processed with CeleScope to remove low quality reads with Cutadapt (v1.17) to trim poly-A tail and adapter sequences. Cell barcode and UMI were extracted. After that, we used STAR (v2.6.1a) ^56^ to map reads to the reference genome GRCh38 (ensembl version 92 annotation). UMI counts and gene counts of each cell were acquired with featureCounts (v2.0.1) ^57^ software, and used to generate expression matrix files for subsequent analysis.

#### Quality control, dimensionality reduction, and batch correction

Single-cell RNA-seq data were processed using Seurat (v5.0.1) ^58^. Quality control was first performed to remove low-quality cells (those with >10% mitochondrial gene content or <300/>3,000 detected genes) and genes expressed in fewer than 10 cells. The filtered data were normalized using the NormalizeData function, followed by cell cycle phase assignment (CellCycleScoring) based on canonical markers. SCTransform normalization was then applied while regressing out technical covariates (mitochondrial content [mitoRatio] and cell cycle scores [S.Score, G2M.Score]). The top 3,000 highly variable genes were selected (FindVariableFeatures) for downstream analysis. Due to the large dataset size (>100,000 cells), batch effects were corrected using reciprocal PCA (RPCA) integration instead of canonical correlation analysis (CCA), as RPCA is more efficient for large-scale datasets.

#### Cell clustering and annotation

Initial cell clustering was performed using the FindClusters function at a resolution of 0.8, identifying 30 distinct clusters. Cell identity was first annotated by mapping our dataset to a publicly available PBMC reference (https://atlas.fredhutch.org/data/nygc/multimodal/pbmc_multimodal.h5seurat) using the MapQuery function ^59^. To identify all the cell clusters in the whole blood sample, we implemented a refined annotation strategy combining reference mapping with canonical marker gene expression: neutrophils were identified by FCGR3B, CXCR2 and G0S2 expression; platelet identity was confirmed using GP9, PF4 and PPBP; while poorly defined “Other” and “OtherT” subpopulations from the reference were excluded. This approach yielded eight well-defined cell populations: neutrophils (FCGR3B^+^, CXCR2^+^, G0S2^+^), monocytes (CD14^+^, PLBD1^+^, PADI4^+^), dendritic cells (CD1C^+^, CST3^+^), B lymphocytes (IGHD^+^, CD79A^+^), CD4^+^ T cells (CD3D^+^, CD3E^+^, CD8A^-^, CD8B^-^), CD8^+^ T cells (CD3D^+^, CD3E^+^, CD8A^+^, CD8B^+^), natural killer cells (KLRB1^+^, KLRD1^+^, FCGR3A^+^), and platelets (GP9^+^, PF4^+^).

Following initial clustering, we subset neutrophils, monocytes, dendritic cells, CD4^+^ T cells, CD8^+^ T cells, NK cells, and B cells as individual Seurat objects for higher-resolution analysis. Each subset underwent reprocessing using standard Seurat workflows: normalization (NormalizeData), feature scaling (ScaleData), variable feature selection (FindVariableFeatures), and dimensionality reduction (RunPCA). Subclusters were then annotated through an integrative approach combining reference dataset projection and canonical marker expression patterns.

#### Differential gene expression analysis

To identify transcriptomic changes associated with LTHAE in each cell types, we performed FindMarkers function. For each defined cell subtype, we employed a Wilcoxon rank-sum test with significance thresholds set at |log2FC| ≥ 0.1 and p-value < 0.05, with complete DEGs lists available in Supplementary Table 5.

To identify conserved transcriptional responses across cell populations, we systematically analyzed the seven major immune cell subtypes (excluding platelets due to their limited transcriptome) and established criteria whereby genes showing significant upregulation (log2FC ≥ 0.1, p-value < 0.05) in at least five of the seven subtypes were classified as consistently upregulated, while those exhibiting significant downregulation (log2FC ≤ -0.1, p-value < 0.05) in at least five subtypes were designated as consistently downregulated.

#### Functional enrichment analysis

Comprehensive functional annotation was performed through Gene Ontology (GO) and Kyoto Encyclopedia of Genes and Genomes (KEGG) pathway enrichment analyses using the clusterProfiler package (v4.10.0) ^60^.

#### Developmental trajectory inference

Pseudotime analysis was performed using Monocle3 (v1.3.4) ^61^ to reconstruct neutrophil differentiation trajectories. Dimensionality reduction was conducted through PCA (50 dimensions) followed by UMAP projection, where the UMAP coordinates were aligned with the pre-computed Seurat embeddings. Cells were clustered using graph-based algorithm before trajectory learning. The pseudotemporal ordering was initiated by manually selecting root nodes (the immature neutrophil cluster G0).

#### CellChat analysis

We used CellChat (v1.6.1) together with its mouse-specific interaction database (CellChatDB.human) to compute communication probabilities for all ligand-receptor pairs under two experimental conditions (plain and LTHAE). Differential communication patterns between plain and long-term high-altitude groups were quantitatively assessed using the rankNet function.

### Integrated transcriptomic-metabolomic pathway analysis

We performed integrative pathway analysis using the joint pathway analysis module on the MetaboAnalyst platform (https://www.metaboanalyst.ca/), which is specifically designed for combined analysis of transcriptomic and metabolomic data at the pathway level. For this analysis, we input both consistently upregulated and downregulated genes and differentially abundant metabolites, together with their respective fold changes. The fold changes of consistently regulated genes were calculated by averaging values across multiple cell subtypes. In the parameter settings, we selected “Metabolic pathways (integrated)” as the pathway database.

## Statistics and reproducibility

Statistical analysis was performed using R software (The R Foundation for Statistical Computing, Vienna, Austria; version 4.1.1). Significance was calculated using two-tailed, unpaired Wilcoxon rank-sum tests unless stated otherwise. Differences were considered statistically significant at the P < 0.05 level (*p < 0.05, **p < 0.01, ***p < 0.001, ****p < 0.001, ns: not significant).

## Supplementary Material

Additional file1: Supplementary Table 1

Volunteer Information and Physiological Data.

Additional file2: Supplementary Table 2, 4

Supplementary Table 2. Effects of high-altitude on physiological parameters in response to LTHAE.

Supplementary Table 4. Differential abundant metabolites with immune-regulatory roles.

Additional file3: Supplementary Table 3

Metabolome Data.

Additional file4: Supplementary Table 5

Different Expression Genes in PBMC Subsets.

Additional file5: Supplementary Figure

## Acknowledgements

We sincerely thank the 46 volunteers who enthusiastically participated in this work. We acknowledge the support from the General Hospital of Tibet Military Area Command.

## Author contributions statement

Y.T.D. and Y.R. designed the project. J.Q.W., W.B.Y., F.S.W., J.J.W and R.Y. performed the bioinformatics analysis. Y.T.D., W.B.Y., C.Z., J.L.Z., Y.H., Y.K.Z, M.Y., M.Y.G., Y.P.T., and Y.R. performed the experiments. Y.T.D., R.Y.X. and J.Q.W. wrote the manuscript. J.L.Z, R.J. and Y.H. conceived and supervised the study. Y.X.X., Y.T.D., Y.R., R.Y.X., J.Q.W. and R.J. contributed to the analysis and interpretation of data.

## Data Availability Statement

The single-cell RNA sequencing (scRNA-seq) and metabolomics data generated in this study have been deposited in the China National GeneBank (CNGB) Sequence Archive (CNSA; https://db.cngb.org/cnsa/) under accession numbers CNP0007414 (scRNA-seq) and CNP0007256 (metabolomics).

All original analysis code has been made freely accessible at https://github.com/Jack123-Wang/High-Altitude-Exposure-Ruan.

## References

1. Su, R. et al. The effects of long-term high-altitude exposure on cognition: A meta-analysis. Neurosci Biobehav Rev 161, 105682 (2024).

2. Richalet, J.P., Hermand, E. & Lhuissier, F.J. Cardiovascular physiology and pathophysiology at high altitude. Nat Rev Cardiol 21, 75–88 (2024).

3. Alcantara-Zapata, D.E., Llanos, A.J. & Nazzal, C. High altitude exposure affects male reproductive parameters: could it also affect the prostate?dagger. Biol Reprod 106, 385–396 (2022).

4. Mallet, R.T., Burtscher, J., Richalet, J.P., Millet, G.P. & Burtscher, M. Impact of High Altitude on Cardiovascular Health: Current Perspectives. Vasc Health Risk Manag 17, 317–335 (2021).

5. Williams, A.M., Levine, B.D. & Stembridge, M. A change of heart: Mechanisms of cardiac adaptation to acute and chronic hypoxia. J Physiol 600, 4089–4104 (2022).

6. Klein, M. et al. Absence of neocytolysis in humans returning from a 3-week high-altitude sojourn. Acta Physiol (Oxf) 232, e13647 (2021).

7. Simpson, L.L., Stembridge, M., Siebenmann, C., Moore, J.P. & Lawley, J.S. Mechanisms underpinning sympathoexcitation in hypoxia. J Physiol 602, 5485–5503 (2024).

8. Zhou, X. et al. Life destiny of erythrocyte in high altitude erythrocytosis: mechanisms underlying the progression from physiological (moderate) to pathological (excessive) high-altitude erythrocytosis. Front Genet 16, 1528935 (2025).

9. Song, Z. et al. Prevalence of High-Altitude Polycythemia and Hyperuricemia and Risk Factors for Hyperuricemia in High-Altitude Immigrants. High Alt Med Biol 24, 132–138 (2023).

10. Yang, X., Liu, H. & Wu, X. High-altitude pulmonary hypertension: a comprehensive review of mechanisms and management. Clin Exp Med 25, 79 (2025).

11. West, J.B. High-altitude medicine. Am J Respir Crit Care Med 186, 1229–1237 (2012).

12. Wang, H. et al. Clinicopathological characteristics of high-altitude polycythemia-related kidney disease in Tibetan inhabitants. Kidney Int 102, 196–206 (2022).

13. Kogut, M.H., Lee, A. & Santin, E. Microbiome and pathogen interaction with the immune system. Poult Sci 99, 1906–1913 (2020).

14. Ganeshan, K. & Chawla, A. Metabolic regulation of immune responses. Annu Rev Immunol 32, 609–634 (2014).

15. Siques, P., Brito, J. & Pena, E. Reactive Oxygen Species and Pulmonary Vasculature During Hypobaric Hypoxia. Front Physiol 9, 865 (2018).

16. Gatterer, H. et al. Altitude illnesses. Nat Rev Dis Primers 10, 43 (2024).

17. Horscroft, J.A. et al. Metabolic basis to Sherpa altitude adaptation. Proc Natl Acad Sci U S A 114, 6382–6387 (2017).

18. Yin, J. et al. Multi-omics reveals immune response and metabolic profiles during high-altitude mountaineering. Cell Rep 44, 115134 (2025).

19. Pham, K., Vargas, A., Frost, S., Shah, S. & Heinrich, E.C. Changes in immune cell populations during acclimatization to high altitude. Physiol Rep 12, e70024 (2024).

20. Shah, Y.M. & Xie, L. Hypoxia-inducible factors link iron homeostasis and erythropoiesis. Gastroenterology 146, 630–642 (2014).

21. Nishide, M. et al. Single-cell multi-omics analysis identifies two distinct phenotypes of newly-onset microscopic polyangiitis. Nat Commun 14, 5789 (2023).

22. Alharthi, S.B. et al. Comparative Study of Complete Blood Count Between High-Altitude and Sea-Level Residents in West Saudi Arabia. Cureus 15, e44889 (2023).

23. Hartmann, S., Krafft, A., Huch, R. & Breymann, C. Effect of altitude on thrombopoietin and the platelet count in healthy volunteers. Thromb Haemost 93, 115–117 (2005).

24. Liew, P.X. & Kubes, P. The Neutrophil’s Role During Health and Disease. Physiol Rev 99, 1223–1248 (2019).

25. Ha, H., Debnath, B. & Neamati, N. Role of the CXCL8-CXCR1/2 Axis in Cancer and Inflammatory Diseases. Theranostics 7, 1543–1588 (2017).

26. Chen, S., Zhu, H. & Jounaidi, Y. Comprehensive snapshots of natural killer cells functions, signaling, molecular mechanisms and clinical utilization. Signal Transduct Target Ther 9, 302 (2024).

27. Ning, Z., Liu, K. & Xiong, H. Roles of BTLA in Immunity and Immune Disorders. Front Immunol 12, 654960 (2021).

28. Mueller, S.N., Gebhardt, T., Carbone, F.R. & Heath, W.R. Memory T cell subsets, migration patterns, and tissue residence. Annu Rev Immunol 31, 137–161 (2013).

29. Sakurai-Yageta, M. & Suzuki, Y. Molecular Mechanisms of Biotin in Modulating Inflammatory Diseases. Nutrients 16 (2024).

30. Rodak, K., Kokot, I. & Kratz, E.M. Caffeine as a Factor Influencing the Functioning of the Human Body-Friend or Foe? Nutrients 13 (2021).

31. Wirthgen, E., Hoeflich, A., Rebl, A. & Gunther, J. Kynurenic Acid: The Janus-Faced Role of an Immunomodulatory Tryptophan Metabolite and Its Link to Pathological Conditions. Front Immunol 8, 1957 (2017).

32. Das, R.K., Brar, S.K. & Verma, M. Recent advances in the biomedical applications of fumaric acid and its ester derivatives: The multifaceted alternative therapeutics. Pharmacol Rep 68, 404–414 (2016).

33. Chen, W., Kuolee, R. & Yan, H. The potential of 3’,5’-cyclic diguanylic acid (c-di-GMP) as an effective vaccine adjuvant. Vaccine 28, 3080–3085 (2010).

34. Gasmi, A. et al. Natural Ingredients to Improve Immunity. Pharmaceuticals (Basel) 16 (2023).

35. Lyon, P., Strippoli, V., Fang, B. & Cimmino, L. B Vitamins and One-Carbon Metabolism: Implications in Human Health and Disease. Nutrients 12 (2020).

36. Liao, W.T. et al. Metabolite Modulation in Human Plasma in the Early Phase of Acclimatization to Hypobaric Hypoxia. Sci Rep 6, 22589 (2016).

37. Gao, J. et al. Metabolomic analysis of human plasma sample after exposed to high altitude and return to sea level. PLoS One 18, e0282301 (2023).

38. Liu, G. et al. Energy metabolic mechanisms for high altitude sickness: Downregulation of glycolysis and upregulation of the lactic acid/amino acid-pyruvate-TCA pathways and fatty acid oxidation. Sci Total Environ 894, 164998 (2023).

39. Burtscher, J. et al. Immune consequences of exercise in hypoxia: A narrative review. J Sport Health Sci 13, 297–310 (2024).

40. Choudhary, R. et al. Respiratory tract infection: an unfamiliar risk factor in high-altitude pulmonary edema. Brief Funct Genomics 23, 38–45 (2024).

41. Lodge, K.M., Cowburn, A.S., Li, W. & Condliffe, A.M. The Impact of Hypoxia on Neutrophil Degranulation and Consequences for the Host. Int J Mol Sci 21 (2020).

42. Gomez, J.C. & Doerschuk, C.M. Hypoxia Can Make Neutrophils Hyper, Potentially Wreaking Havoc during Exacerbations in Chronic Obstructive Pulmonary Disease. Am J Respir Crit Care Med 205, 862–864 (2022).

43. Yao, S. et al. The association between altitude and serum folate levels in Tibetan adults on the Tibetan plateau. Sci Rep 12, 17886 (2022).

44. Gombart, A.F., Pierre, A. & Maggini, S. A Review of Micronutrients and the Immune System-Working in Harmony to Reduce the Risk of Infection. Nutrients 12 (2020).

45. Newstead, S. Structural basis for recognition and transport of folic acid in mammalian cells. Curr Opin Struct Biol 74, 102353 (2022).

46. Kapellos, T.S. et al. Human Monocyte Subsets and Phenotypes in Major Chronic Inflammatory Diseases. Front Immunol 10, 2035 (2019).

47. Macri, C., Pang, E.S., Patton, T. & O’Keeffe, M. Dendritic cell subsets. Semin Cell Dev Biol 84, 11–21 (2018).

48. Leylek, R. et al. Integrated Cross-Species Analysis Identifies a Conserved Transitional Dendritic Cell Population. Cell Rep 29, 3736–3750 e3738 (2019).

49. Nutt, S.L., Hodgkin, P.D., Tarlinton, D.M. & Corcoran, L.M. The generation of antibody-secreting plasma cells. Nat Rev Immunol 15, 160–171 (2015).

50. Chung, J.B., Silverman, M. & Monroe, J.G. Transitional B cells: step by step towards immune competence. Trends Immunol 24, 343–349 (2003).

51. Lam, J.H., Smith, F.L. & Baumgarth, N. B Cell Activation and Response Regulation During Viral Infections. Viral Immunol 33, 294–306 (2020).

52. Akkaya, M., Kwak, K. & Pierce, S.K. B cell memory: building two walls of protection against pathogens. Nat Rev Immunol 20, 229–238 (2020).

53. Li, J. et al. B cell metabolism in autoimmune diseases: signaling pathways and interventions. Front Immunol 14, 1232820 (2023).

54. Sun, L., Su, Y., Jiao, A., Wang, X. & Zhang, B. T cells in health and disease. Signal Transduct Target Ther 8, 235 (2023).

55. Dura, B. et al. scFTD-seq: freeze-thaw lysis based, portable approach toward highly distributed single-cell 3’ mRNA profiling. Nucleic Acids Res 47, e16 (2019).

56. Dobin, A. et al. STAR: ultrafast universal RNA-seq aligner. Bioinformatics 29, 15–21 (2013).

57. Liao, Y., Smyth, G.K. & Shi, W. featureCounts: an efficient general purpose program for assigning sequence reads to genomic features. Bioinformatics 30, 923–930 (2014).

58. Hao, Y. et al. Dictionary learning for integrative, multimodal and scalable single-cell analysis. Nat Biotechnol 42, 293–304 (2024).

59. Hao, Y. et al. Integrated analysis of multimodal single-cell data. Cell 184, 3573–3587 e3529 (2021).

60. Wu, T. et al. clusterProfiler 4.0: A universal enrichment tool for interpreting omics data. Innovation (Camb) 2, 100141 (2021).

61. Cao, J. et al. The single-cell transcriptional landscape of mammalian organogenesis. Nature 566, 496–502 (2019).

