## Supplementray Figure for "Remodeling of the immune-metabolic landscape triggered by long-term high-altitude exposure"

**Supplementary Figure Legends**

**Supplementary Figure 1. Cellular and functional heterogeneity in the T cell under long-term high-altitude exposure**

A. UMAP plot of the CD8+ T cell subtypes.

B. Dot plots depicting the percentages and average expressions of marker genes in CD8+ T cell subtypes.

C. Cell density plots of CD8+ T cell in LA and HA groups, with color intensity proportional to local cell density.

D. Violin plots showing the changes in the relative proportions of CD8+ T cell subtypes between LA and HA groups.

E. Multi-group volcano plots displaying DEGs across CD8+ T cell subtypes in LA and HA groups, with significantly upregulated (red) and downregulated (blue) genes indicated.

F. Bar plot depicting the number of DEGs in each CD8+ T cell subtype between LA and HA groups.

G. Functional enrichment analysis of DEG between LA and HA groups across CD8+ T cell subtypes. The bubble plots visualize significantly enriched GO terms, with color gradients representing -log2(P-values) for upregulated (red) and downregulated (blue) gene sets, and bubble size corresponds to the number of enriched genes.

H. UMAP plot of the CD4+ T cell subtypes.

I. Dot plots depicting the percentages and average expressions of marker genes in CD4+ T cell subtypes.

J. Cell density plots of CD4+ T cell in LA and HA groups, with color intensity proportional to local cell density.

K. Violin plots showing the changes in the relative proportions of CD4+ T cell subtypes between LA and HA groups.

L. Multi-group volcano plots displaying DEGs across CD4+ T cell subtypes in LA and HA groups, with significantly upregulated (red) and downregulated (blue) genes indicated.

M. Bar plot depicting the number of DEGs in each CD4+ T cell subtype between LA and HA groups.

N. Functional enrichment analysis of DEG between LA and HA groups across CD4+ T cell subtypes.

**Figure S1**

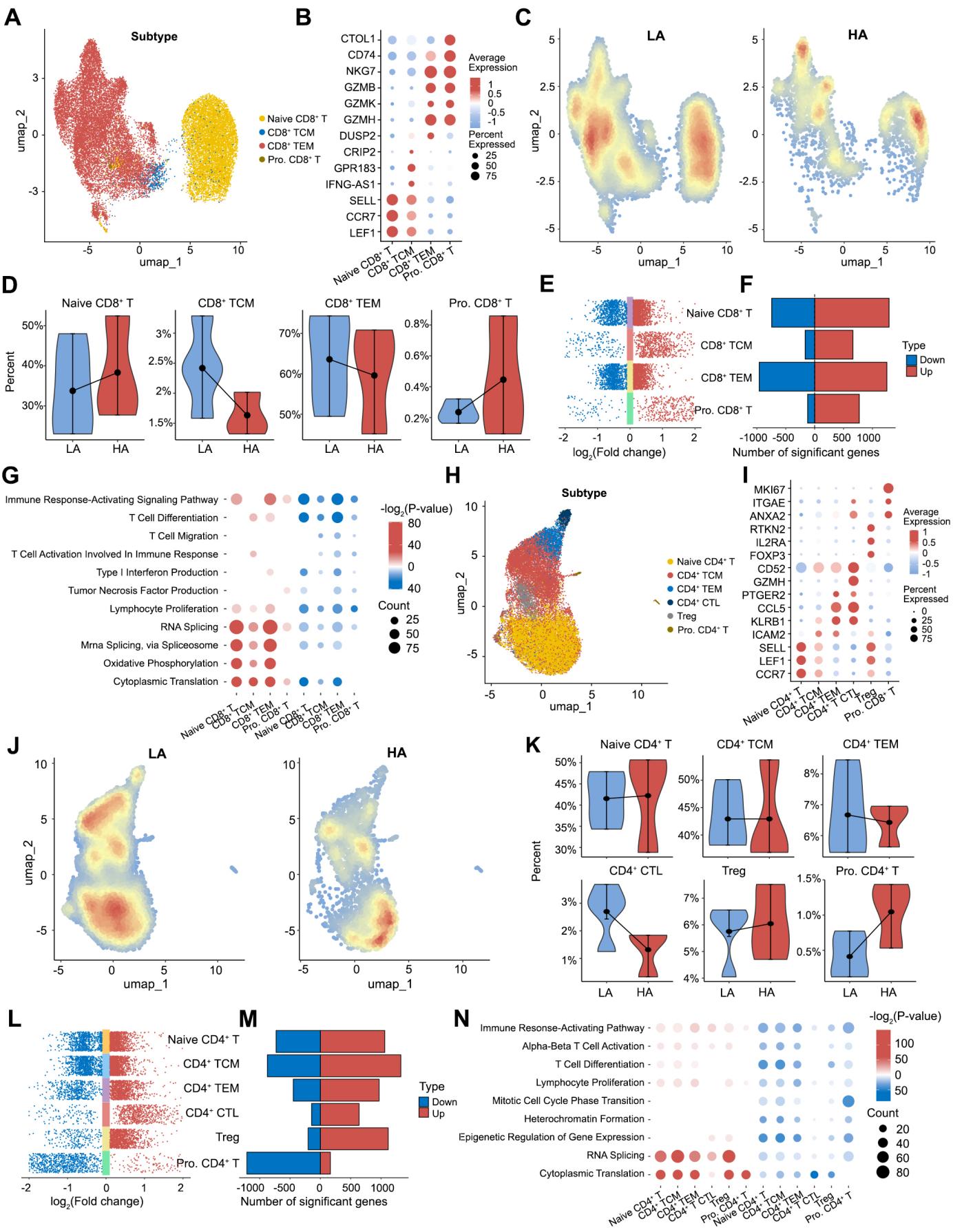
