## Supplementray Table2,4 for "Remodeling of the immune-metabolic landscape triggered by long-term high-altitude exposure"

**List of supplementary information**

**Supplementary Table 2.** Effects of high-altitude on physiological parameters in response to long-term high-altitude exposure.

**Supplementary Table 4.** Differential abundant metabolites with immune-regulatory roles.

**Supplementary Table 2.** Effects of high-altitude on physiological parameters in response to long-term high-altitude exposure.

| **Variables** | **Low-altitude** | **High-altitude** |
| --- | --- | --- |
| Gender (male/female, n) | 45/0 | 40/0 |
| Age (years) | 22.0±1.6 | 22.1±1.7 |
| Height (cm) | 176.2±3.8 | 176.1±4.0 |
| Body weight (kg) | 70.5±5.3 | 66.5±5.1 |
| Systolic blood pressure (mmHg) | 116.0±10.5 | 105.8±11.9 |
| Diastolic blood pressure (mmHg) | 62.6±6.6 | 65.3±8.3 |
| Blood oxygen saturation (%) | 99.2±0.5 | 86.7±2.5 |
| Body mass index (kg/m^2^) | 22.7±1.5 | 21.4±1.4 |
| Heart rate (beats per minute) | 74.2±9.8 | 91.0±14.4 |
| Blood routine |  |  |
| White blood cell count (×10^9^/L) | 6.0±1.2 | 7.0±1.6 |
| Red blood cell count (×10^12^/L) | 4.9±0.3 | 5.8±0.3 |
| Hemoglobin (g/L) | 158.0±54.4 | 172.0±8.3 |
| Platelet count(×10^9^/L) | 252.9±46.1 | 273.8±48.7 |
| Blood chemistry |  |  |
| Triglyceride(mmol/L) | 0.8±0.3 | 0.6±0.2 |
| Total cholesterol(mmol/L) | 4.0±0.7 | 4.1±0.6 |
| High-density lipoprotein(mmol/L) | 1.3±0.2 | 1.3±0.3 |
| Low-density lipoprotein(mmol/L) | 2.4±0.5 | 2.0±0.4 |
| Creatine kinase(U/L) | 203.6±248.7 | 1692.0±5863.0 |
| Creatine kinase isoenzyme(U/L) | 12.6±5.5 | 38.7±56.4 |
| Cell count |  |  |
| Neu (×10^9^/L) | 3.6±1.2 | 4.0±1.4 |
| Lymph (×10^9^/L) | 1.9±0.5 | 2.3±0.5 |
| Mon (×10^9^/L) | 0.3±0.09 | 0.5±0.2 |
| Eos (×10^9^/L) | 0.15±0.08 | 0.13±0.11 |
| Bas (×10^9^/L) | 0.03±0.02 | 0.01±0.01 |
| Cell percentage (%) |  |  |
| Neu (%) | 58.8±9.8 | 56.2±8.9 |
| Lymph (%) | 32.5±8.9 | 34.0±8.5 |
| Mon (%) | 5.8±1.2 | 7.8±2.4 |
| Eos (%) | 2.6±1.6 | 1.8±1.3 |
| Bas (%) | 0.5±0.3 | 0.1±0.1 |

**Supplementary Table 4.** Differential abundant metabolites with immune-regulatory roles.

| Type | Metabolites | Function | Mechanism |
| --- | --- | --- | --- |
| Up-regulated | Biotin | Anti-inflammatory | Inhibits the production of pro-inflammatory factors including TNF-α, IL-1β, IFN-γ, and IL-17^1^ |
|  | Caffeine | Anti-inflammatory | Reduces the levels of CRP and inflammatory cytokines such as IL-1β, IL-6, IL-18, and TNF-α, and promotes the secretion of anti-inflammatory factor IL-10^2^ |
|  | Theobromine | Anti-inflammatory | Suppresses B cell activation and antibody production while impairing T cell maturation^3^ |
|  | Inosine | Anti-inflammatory | Suppresses T cell receptor signaling, decreases TNF-α and IL-6 production, and inhibits immune cell proliferation^4^ |
|  | Rauwolscine | Anti-inflammatory | Suppresses macrophage activation and inhibits inflammatory cell infiltration^5^ |
|  | Eudesmin | Anti-inflammatory | Suppresses T cell proliferation and inhibits mast cell histamine release^6^ |
|  | Kynurenic acid | Anti-inflammatory | Exerts antioxidant effects by scavenging free radicals, suppresses pro-inflammatory cytokine secretion, and inhibits T-cell proliferation^7^ |
|  | N-Acetylserotonin | Anti-inflammatory | Suppresses TNF-α production and scavenges reactive oxygen species (ROS)^8^ |
|  | Smilagenin | Anti-inflammatory | Inhibits both nitric oxide (NO) production and NF-κB expression^9^ |
|  | Diacetyl | Anti-inflammatory | Suppresses NLRP3 inflammasome activation via NF-κB pathway modulation^10^ |
|  | Docosahexaenoic acid | Anti-inflammatory | Suppresses both MAPK/NF-κB signaling and apoptotic pathways^11^ |
|  | Fumaric acid | Anti-inflammatory | Suppresses secretion of pro-inflammatory cytokines (TNF-α, IL-6, IL-23) and inhibits activation of dendritic cells, macrophages, and T lymphocytes^12^ |
| Type | **Metabolites** | **Function** | **Mechanism** |
| Down-regulated | 3',5'-Cyclic GMP | Pro-inflammatory | Promotes inflammatory cytokine and chemokine secretion, activates dendritic cell maturation, and promotes Th1-type immune polarization^13^ |
|  | Cholic acid | Anti-inflammatory | Promotes regulatory T cell (Treg) differentiation, induces anti-inflammatory dendritic cell (DC) polarization, enhances immune tolerance, and shifts cytokine balance toward anti-inflammatory mediators while suppressing pro-inflammatory factors^14^ |
|  | Ascorbic acid | Anti-inflammatory | Inhibits pro-inflammatory cytokines and reduces histamine levels^15^ |
|  | Folic acid | Pro-inflammatory | Promotes antiviral and innate immune pro-inflammatory responses^16^ |
|  | Coptisine | Anti-inflammatory | Exerts multifaceted therapeutic effects, including anti-inflammatory, immunomodulatory, antioxidant, and anti-fibrotic activities, primarily through modulation of key signaling pathways such as PI3K/AKT, Th17 cell differentiation, and inflammatory bowel disease-related pathways^17^ |

**References**

1.	Sakurai-Yageta, M. & Suzuki, Y. Molecular Mechanisms of Biotin in Modulating Inflammatory Diseases. *Nutrients* **16** (2024).

2.	Rodak, K., Kokot, I. & Kratz, E.M. Caffeine as a Factor Influencing the Functioning of the Human Body-Friend or Foe? *Nutrients* **13** (2021).

3.	Camps-Bossacoma, M., Perez-Cano, F.J., Franch, A. & Castell, M. Theobromine Is Responsible for the Effects of Cocoa on the Antibody Immune Status of Rats. *J Nutr* **148**, 464-471 (2018).

4.	Samami, E. *et al.* Inosine, gut microbiota, and cancer immunometabolism. *Am J Physiol Endocrinol Metab* **324**, E1-E8 (2023).

5.	Kambhampati, V., Eedara, A. & Andugulapati, S.B. Yohimbine treatment improves pulmonary fibrosis by attenuating the inflammation and oxidative stress via modulating the MAPK pathway. *Biochem Pharmacol* **230**, 116613 (2024).

6.	Hong, P.T.L., Kim, H.J., Kim, W.K. & Nam, J.H. Flos magnoliae constituent fargesin has an anti-allergic effect via ORAI1 channel inhibition. *Korean J Physiol Pharmacol* **25**, 251-258 (2021).

7.	Wirthgen, E., Hoeflich, A., Rebl, A. & Gunther, J. Kynurenic Acid: The Janus-Faced Role of an Immunomodulatory Tryptophan Metabolite and Its Link to Pathological Conditions. *Front Immunol* **8**, 1957 (2017).

8.	Perianayagam, M.C., Oxenkrug, G.F. & Jaber, B.L. Immune-modulating effects of melatonin, N-acetylserotonin, and N-acetyldopamine. *Ann N Y Acad Sci* **1053**, 386-393 (2005).

9.	Cortes, A.J. *et al.* Steroidal saponin from Agave marmorata Roezl modulates inflammatory response by inhibiting NF-kappaB and AP-1. *Nat Prod Res* **36**, 1123-1128 (2022).

10.	Park, M.H. *et al.* N,N'-Diacetyl-p-phenylenediamine restores microglial phagocytosis and improves cognitive defects in Alzheimer's disease transgenic mice. *Proc Natl Acad Sci U S A* **116**, 23426-23436 (2019).

11.	Jiang, Z. *et al.* Lycium barbarum glycopeptide alleviates neuroinflammation in spinal cord injury via modulating docosahexaenoic acid to inhibiting MAPKs/NF-kB and pyroptosis pathways. *J Transl Med* **21**, 770 (2023).

12.	Das, R.K., Brar, S.K. & Verma, M. Recent advances in the biomedical applications of fumaric acid and its ester derivatives: The multifaceted alternative therapeutics. *Pharmacol Rep* **68**, 404-414 (2016).

13.	Chen, W., Kuolee, R. & Yan, H. The potential of 3',5'-cyclic diguanylic acid (c-di-GMP) as an effective vaccine adjuvant. *Vaccine* **28**, 3080-3085 (2010).

14.	Su, X., Gao, Y. & Yang, R. Gut microbiota derived bile acid metabolites maintain the homeostasis of gut and systemic immunity. *Front Immunol* **14**, 1127743 (2023).

15.	Carr, A.C. & Maggini, S. Vitamin C and Immune Function. *Nutrients* **9** (2017).

16.	Gasmi, A. *et al.* Natural Ingredients to Improve Immunity. *Pharmaceuticals (Basel)* **16** (2023).

17.	Yang, Y. *et al.* Therapeutic targets and pharmacological mechanisms of Coptidis Rhizoma against ulcerative colitis: Findings of system pharmacology and bioinformatics analysis. *Front Pharmacol* **13**, 1037856 (2022).
